# Attenuating the DNA damage response to double strand breaks restores function in models of CNS neurodegeneration

**DOI:** 10.1101/440636

**Authors:** Richard I. Tuxworth, Matthew J. Taylor, Ane Martin Anduaga, Alaa Hussien-Ali, Sotiroula Chatzimatthaiou, Joanne Longland, Adam M. Thompson, Sharif Almutiri, Pavlos Alifragis, Charalambos P. Kyriacou, Boris Kysela, Zubair Ahmed

## Abstract

DNA double-strand breaks are a feature of many acute and long-term neurological disorders, including neurodegeneration, following neurotrauma and after stroke. Persistent activation of the DNA damage response in response to double strand breaks contributes to neural dysfunction and pathology as it can force post-mitotic neurons to re-enter the cell cycle leading to senescence or apoptosis. Mature, non-dividing neurons may tolerate low levels of DNA damage, in which case muting the DNA damage response might be neuroprotective. Here, we show that attenuating the DNA damage response by targeting the meiotic recombination 11, Rad50, Nijmegen breakage syndrome 1 complex, which is involved in double strand break recognition, is neuroprotective in three neurodegeneration models in *Drosophila* and prevents Aβ_1-42_-induced loss of synapses in embryonic hippocampal neurons. Attenuating the DNA damage response after optic nerve injury is also neuroprotective to retinal ganglion cells and promotes dramatic regeneration of their neurites both *in vitro* and *in vivo*. Dorsal root ganglion neurons similarly regenerate when the DNA damage response is targeted *in vitro* and *in vivo* and this strategy also induces significant restoration of lost function after spinal cord injury. We conclude that muting the DNA damage response in the nervous system is neuroprotective in multiple neurological disorders. Our results point to new therapies to maintain or repair the nervous system.

## Introduction

DNA double-strand breaks are the most deleterious type of DNA damage. In mitotically cycling cells, double strand breaks trigger the DNA damage response to arrest the cell cycle in and mount repair via non-homologous end-joining in G1 or G2 phases or homologous recombination in M and S phases (Mladenov *et al.*, 2016). Double strand breaks are a feature of many acute and long-term neurological disorders, including many forms of neurodegeneration (Simpson *et al.*, 2015; Merlo *et al.*, 2016), following neurotrauma (Kotipatruni *et al.*, 2011) and after stroke (Hayashi *et al.*, 1998). Unrepaired double strand breaks in neurons leads to persistent activation of the DNA damage response which, in turn, is a trigger for dysregulation of the cell cycle and aberrant re-entry of neurons into G1 leading to neural dysfunction, apoptosis and senescence (Herrup *et al.*, 2004). Re-entry of neurons into the cell cycle and senescence are features of Alzheimer’s disease (McShea *et al.*, 1997; Nagy *et al.*, 1997; Fielder *et al.*, 2017) whilst dysfunctional regulation of the cell cycle and has also been documented in models of cerebral ischemia (Katchanov *et al.*, 2001; Wen *et al.*, 2004) and in post-mortem brain samples of stroke patients (Love, 2003).

If persistent activation of the DNA damage response is a trigger for neuronal dysfunction, apoptosis and senescence then potentially muting the DNA damage response would be neuroprotective. A key sensor and early processor of double strand breaks is the MRN complex, comprising the Mre11, Rad50 and Nbs1/Nbn proteins (Lamarche *et al.*, 2010). Association of the MRN complex to double strand breaks leads to recruitment and activation of the ataxia telangiectasia mutated (ATM) kinase, which coordinates multiple arms of the DNA damage response, including cell-cycle arrest, repair and apoptosis (Shiloh and Ziv, 2013). ATM activation is particularly associated with non-homologous end-joining; in contrast, the related kinase, ataxia telangiectasia and Rad3-related (ATR) protein, regulates homologous recombination. Double strand breaks are characterised by the activation of sensor kinases, including DNA-protein kinase catalytic subunit and regions of DNA damage incorporating phosphorylated histone γH2Ax (Shrivastav *et al.*, 2008). Therefore, γH2Ax is commonly used to monitor direct activation of the DNA damage response in terms of double strand breaks (Celeste *et al.*, 2002; Simpson *et al.*, 2015; Milanese *et al.*, 2018). ATM is recruited at the site of lesions such as double strand breaks to phosphorylate in cis the histone H2Ax (γH2Ax) (Celeste *et al.*, 2002). γH2Ax marks the initiation of a nucleation process resulting in the formation of characteristic γH2Ax^+^ nuclear foci, which leads to additional ATM recruitment at the damage site, thereby escalating the kinase activity (Shiloh and Ziv, 2013).

In post-mitotic neurons, homologous recombination is unlikely to feature in the repair of double strand breaks as no sister chromatid is available to act as a template for repair. Given its position at the apex of the DNA damage response, the MRN complex is a potential target to attenuate or mute the DNA damage response in neurons. Here, we targeted the MRN complex using both genetic approaches in *Drosophila* models of neurodegeneration and using small-molecule inhibitors in two rat models of acute neurotrauma: ocular injury and spinal cord injury. We identified that targeting of the MRN complex was neuroprotective in each scenario. This universality opens up new possibilities for treating human neurological disorders.

## Materials and Methods

### *Drosophila* stocks and breeding

For all *Drosophila* experiments except circadian analysis, expression of the neurodegeneration-associated transgenes was restricted to adult neurons by use of an *elav^C155^*; *GAL80^ts^* driver line. *Drosophila* were bred on standard yeast/agar media in bottles at 18°C on a 12 h light/dark cycle until eclosion. To induce expression, flies were shifted to and maintained at 29°C, 70% humidity on a 12 h light/dark cycle for the duration of the experiment. Virgin females of the driver line were crossed to males of the control or experimental lines. The control used was an isogenic *w^1118^* strain and all experimental lines were backcrossed over 5 generations into this line. UAS-lines and the *rad50^EP1^* allele were followed during backcrossing by the presence of the w^+^ transgene and the *nbs* alleles by PCR. All fly stocks were obtained from the Vienna or Bloomington *Drosophila* stock centres except for UAS-Aβ_1-42_ (12-linker) (Speretta *et al.*, 2012), which was a kind gift of Dr. Damien Crowther (University of Cambridge) and UAS-0N4R Tau R406W (Wittmann *et al.*, 2001), which was a kind gift of Dr. Mel Feany (Harvard University).

### Quantification of fly movement

Newly eclosed adult flies were separated into cohorts of 20 flies in vials and shifted to 29°C to begin expression by climbing assay or movement tracking.

#### Negative geotaxis climbing assays

To test climbing, flies were removed from the incubator and allowed to acclimatise for 1 h to the room temperature and humidity. Each vial of flies was tipped into an empty vial and left for 60 s to acclimatise. Flies were tapped to the base and allowed to climb up for 30 s. The percentage of flies climbing above a line 2.5 cm above the base was recorded as the mean of three repeats. Groups of 5 vials were tested together. Flies were transferred to fresh food vials after testing and returned to the incubator. For the Htt.Q128 experiment, second order polynomials were fitted to the data by non-linear regression in Prism 7 and compared by sum-of-squares F-test. For the Aβ_1-42_ experiment, data were compared by 2-way ANOVA. Significance in both cases was set at p<0.05.

#### Movement tracking with DART

Flies were housed individually in 65 × 5 mm locomotor tubes (Trikinetics). Tubes were mounted in groups of 20 horizontally on custom platforms and movement recorded from above via a Logitech C920 HD camera mounted on a copy stand. The position of each fly was determined at 5 Hz and movement quantified using DART software running in MATLAB 2017a (Faville *et al.*, 2015). Vibrational stimulation was applied 5 times at 10 min intervals to the flies via motors mounted to the underside of each platform and controlled by the DART software. The mean speed of the population of 20 flies over the 120 s before each stimulation, and the maximum amplitude of the population response to stimulation were quantified by DART. Sigmoidal trend lines were fitted to the data by non-linear regression in Prism 7 and compared by sum-of-squares F-test. A full description of the adaptation of the DART circadian behaviour system to quantify the startle response of flies will be published elsewhere.

### Circadian analysis

*tim-Gal4* was used to express Aβ_1-42_ in clock neurons. Flies were maintained at 25°C throughout development and into adulthood in bottles. Newly eclosed adult flies were sorted for genotype and transferred to individual 65 × 5 mm locomotor tubes in DAM activity monitors (Trikinetics) and maintained in 12 h-light/12 h-dark cycle for 3 d then in 12 h-dark/12 h-dark for 10 d. Circadian behaviour was analysed with both spectral analysis using CLEAN and cosinor as described previously (Levine *et al.*, 2002; Rosato and Kyriacou, 2006). Activity in the first day of DD was discarded. Genotypes were compared by ANOVA with a Tukey post-hoc test. Additional flies from the same breeding cohorts were sorted for genotype but maintained in vials in LD to maintain circadian cohesion. After 10 d the brains were dissected at ZT0 (just before lights on) and fixed and stained essentially as described (Dissel *et al.*, 2004) except that the brains were incubated in primary antibodies for 4 d and in secondary antibody overnight (both at 4°C). Primary antibodies used were: mouse anti-PDF C7-s (1:50; Developmental Studies Hybridoma Bank) and rabbit anti-Per1 (1:200; Santa Cruz, CA, USA). Secondary antibodies were: Alexa-488 goat anti-mouse IgG and Cy3 goat anti-rabbit IgG (both 1:100 dilution; ThermoFisher, Leicester, UK). Brains were viewed using an Olympus FV1000 confocal microscope. Per^+^ cells were counted manually and their identities determined based on position. for statistical analysis, the PDF^+^ lLNv and sLNv cells were considered together and, similarly, the PDF^−^ DN1, DN2 and LNd cells were pooled. At least 7 brains were used per genotype. Cell numbers were compared by Kruskal-Wallis with a Dunn’s post-hoc test.

### Drugs

Mirin and KU-60019 were both purchased from Tocris, Bristol, UK. Mirin is a small molecule inhibitor that blocks the 3’ and 5’ exonuclease activity associated with Mre11 and prevents ATM activation in response to double strand breaks (Dupre *et al.*, 2008; Stivers, 2008; Garner *et al.*, 2009). KU-60019 is a potent ATM kinase inhibitor and has been used to inhibit migration and invasion of human glioma cells *in vitro* (Golding *et al.*, 2009).

### Rat primary hippocampal neuron culture

Primary cultures were prepared from dissected hippocampi of E18 Sprague Dawley rat embryos as previously described (Russell *et al.*, 2012). Cells were plated at a density of either 75,000 or 500,000 cells on poly-D-lysine (Sigma, Poole, UK; 0.1 mg/ml in borate buffer, pH 8.5) coated glass cover slips or 6-well plates respectively. The plating medium was DMEM supplemented with 5% FBS, penicillin/streptomycin (P/S) and 0.5 mM L-glutamine (all from ThermoFisher). On the next day, the medium was changed to Neurobasal medium supplemented with B27, P/S and 0.5 mM L-glutamine (ThermoFisher). Cultures were incubated at 37°C and 5% CO_2_ and were used between 18 to 21 days *in vitro*.

### Rat primary DRGN cultures

Primary adult rat DRGN were prepared as described previously (Ahmed *et al.*, 2005). DRGN cells were cultured in Neurobasal-A (ThermoFisher) at a plating density of 500 DRGN/well in chamber slides (Beckton-Dickinson, Watford, UK) pre-coated with 100 μg/ml poly-D-lysine. In preliminary experiments, the optimal concentration of mirin (100 μM) and KU-60019 (10 μM) that promoted DRGN survival and neurite outgrowth were determined. The positive control was pre-optimised fibroblast growth factor (FGF)-2 (Peprotech, London, UK; 10 ng/ml (Ahmed *et al.*, 2005)). Cells were cultured for 4 days in a humidified chamber at 37 °C and 5% CO_2_.

### Rat primary retinal cultures

Primary adult retinal cultures, containing enriched populations of retinal ganglion cells (RGC) were prepared as described previously (Ahmed *et al.*, 2006b). Briefly, retinal cells were dissociated into single cell suspensions using a Papain dissociation kit (Worthington Biochemicals, New Jersey, USA). 125 × 10^3^ retinal cells/well were cultured in chamber slides (Beckton-Dickinson) pre-coated with 100 μg/ml poly-D-lysine. In preliminary experiments, the optimal concentration of mirin (100 μM) and KU-60019 (10 μM) that promoted RGC survival and neurite outgrowth were determined. The positive control was pre-optimised CNTF (Peprotech; 20 ng/ml (Douglas *et al.*, 2009)). Cells were cultured for 4 days in a humidified chamber at 37°C and 5% CO_2_.

### Preparation of A**β** oligomers and neuronal treatment

Preparation of Aβ peptides and treatment of neurons has been previously described (Marsh *et al.*, 2017). Briefly, Aβ_1-42_ (Bachem) was prepared by dissolving the peptide in DMSO (1 mM). The reconstituted peptides were diluted in 10 mM Tris, pH7.8 at a working concentration of 300 μM and stored in aliquots at −80 °C. Peptides were diluted to working concentration in full Neurobasal medium after thawing and used immediately. Neurons were incubated with 0.5 μM Aβ_1-42_ or 0.5 μ Aβ_1-42_ + 100 μM mirin for 24 h.

### Immunocytochemistry

*Adult fly brains* were dissected in PBS and fixed for 20 min in 4% formaldehyde in PBS. After washing, brains were blocked in 1% BSA in PBS for 1 h then incubated with rat anti-Elav (Developmental Studies Hybridoma Bank clone 7E8A1, 1:25) and mouse anti-phosphorylated pH2Av (DSHB clone UNC93-5.2.1, 1:25) for 48 h at 4°C. Brains were washed in PBS + 0.3% triton X-100 for 2 h at RT then incubated in cross-adsorbed Alexa-594 anti-rat and Alexa-488 anti-mouse secondary antibodies (Jackson ImmunoResearch, both 1:400) for 48 h at 4°C. After washing as before, brains were mounted on bridge slides in Vectashield (Vector Labs).

*Hippocampal cells* on coverslips following treatment, were rinsed once with pre-warmed (37 °C) PBS and fixed with 4% paraformaldehyde (PFA; TAAB, Peterborough, UK), 2% Sucrose in PBS. Fixed neurons were washed with TBS, permeabilised with 0.1% Tween-20 and 5% horse serum in TBS for 45 min at room temperature and incubated with primary antibody overnight at 4 °C. Coverslips were mounted using the ProLong Gold reagent (ThermoFisher). The antibodies used were: mouse anti-H2Ax pSer319 (γH2Ax; JBW301; 1;1000 dilution; Merck) and Alexa-568 anti-mouse IgG (ThermoFisher). The cells were visualized on a spin disc confocal system (CARV from Digital Imaging Solutions) with an EM-CCD camera (Rolera/QI Cam 3500) mounted on an Olympus X71 microscope, using a 100x fluoplan objective (NA 4.2). The microscope confocal system was supported by Image Pro 6.0 software.

*DRGN and RGC* were fixed in 4% paraformaldehyde, washed in 3 changes of PBS before being subjected to immunocytochemistry as described previously (Ahmed *et al.*, 2005; Ahmed *et al.*, 2006b). Antibodies used were: mouse anti-βIII tubulin (1:200 dilution; Sigma) to visualise neuronal cell soma and neurites and Alexa-488 goat anti-mouse IgG (1:400 dilution; ThermoFisher). Slides were then viewed with an epifluorescent Axioplan 2 microscope, equipped with an AxioCam HRc and running Axiovision Software (all from Zeiss, Hertfordshire, UK). The proportion of DRGN with neurites and the mean neurite length were calculated using Axiovision Software by an investigator masked to the treatment conditions as previously described (Ahmed *et al.*, 2005; Ahmed *et al.*, 2006b). All experiments were performed in triplicate and repeated on 3 independent occasions.

### *In vivo* surgical procedures

Experiments were licensed by the UK Home Office and all experimental protocols were approved by the University of Birmingham’s Animal Welfare and Ethical Review Board. All animal surgeries were carried out in strict accordance to the guidelines of the UK Animals Scientific Procedures Act, 1986 and the Revised European Directive 1010/63/EU and conformed to the guidelines and recommendation of the use of animals by the Federation of the European Laboratory Animal Science Associations (FELASA). Animals were housed in a standard facility and kept on a 12-h light–12-h dark cycle, with a daytime luminance of 80 lux, fed and watered ad libitum. For all *in vivo* experiments, adult female Sprague-Dawley rats weighing 170-220g (Charles River, Margate, UK) were used. Rats were randomly assigned to each experimental group with the investigators masked to the treatment conditions.

#### Optic nerve crush injury (ONC)

Rats were injected subcutaneously with 50 µl Buprenorphine to provide analgesia prior to surgery and anaesthetised using 5% of Isoflurane in 1.8 ml/l of O_2_ with body temperature and heart rate monitored throughout surgery. ONC was performed 2mm from the lamina cribrosa using watchmaker’s forceps (Ahmed *et al*., 2005; Berry *et al*., 1996).

In pilot dose-finding experiments, mirin and KU-60019 were intravitreally injected at 1, 2, 2.5, 5, 7.5 and 10 µg (n = 3 rats/group, 2 independent repeats), without damaging the lens, immediately after ONC and every other day, or twice weekly or once every 7 days, in a final volume of 5µl saline for 24 days (not shown). Rats were then killed and retinae were dissected out, lysed in ice-cold lysis buffer, separated on 12% SDS PAGE gels and subjected to western blot detection of γH2Ax levels. We determined that the dosing frequency of twice weekly and 2.5 and 5 µg of mirin and KU-60019 optimally reduced γH2Ax levels, respectively. Twice weekly intravitreal injections were sufficient in the eye since the vitreous may acts as a slow release gel due to its composition, being made of mainly water, collagen type II fibrils with glycosaminoglycans, hyaluronan and opticin. Optimal doses were then used for all experiments described in this manuscript. Rats were killed in rising concentrations of CO_2_ at 1 and 24 days after optic nerve crush injury for western blot analyses or at 24 days after ONC for determination of RGC survival and axon regeneration, as described below.

For the experiments reported in this manuscript, n = 6 rats/group were used and assigned to: (1), Intact controls (no surgery to detect baseline parameters); (2), ONC+vehicle (ONC followed by intravitreal injection of vehicle solution; to detect surgery-induced changes); (3), ONC+mirin (ONC followed by intravitreal injection of 2.5µg of mirin, twice weekly; to monitor effects of inhibiting Mre11); and (4), ONC+KU-60019 (ONC followed by intravitreal injection of 5µg of KU-60019; to monitor effects of inhibiting ATM). FluoroGold was injected into the proximal nerve stump 2d before sacrifice and whole retinal flatmounts were used to assess RGC survival *in vivo*, as described by us previously (Ahmed *et al.*, 2011). Each experiment was repeated on 3 independent occasions with a total n = 18 rats/group/test.

#### DC crush injury

For the DC lesion model, experiments also comprised n = 6 rats/group: (1), Sham controls (Sham; to detect surgery-induced changes; partial laminectomy but no DC lesion); (2), DC transected controls + intrathecal injection of vehicle (PBS); (3), (DC+vehicle; to detect injury-mediated changes); (4), DC+intrathecal injection of mirin (DC+mirin; to monitor effects of inhibiting Mre11); (5), and DC+intrathecal injection of KU-60019 (DC+KU-60019; to monitor effects of inhibiting ATM). Each experiment was repeated on 3 independent occasions with a total n = 18 rats/group/test.

Rats were injected subcutaneously with 50 μl Buprenorphine to provide analgesia prior to surgery and anaesthetised using 5% of Isoflurane in 1.8 ml/l of O_2_ with body temperature and heart rate monitored throughout surgery. For optic nerve crush injury, a supraorbital approach was used to expose the optic nerve and crushed with calibrated watchmaker’s forceps 2mm from the lamina cribrosa of the eye (Berry *et al.*, 1996). Animals were immediately injected intravitreally and avoiding damaging the lens with vehicle, mirin or KU-60019. Animals were then allowed to survive for 24 days before assessment of RGC survival and axon regeneration. For the DC injury model, a partial T8 laminectomy was performed and the DC were crushed bilaterally using calibrated watchmaker’s forceps as described (Surey *et al.*, 2014; Almutiri *et al.*, 2018). The subarachnoid space was cannulated with a polyethylene tube (PE-10; Beckton Dickinson) through the atlanto-occipital membrane as described by others (Yaksh and Rudy, 1976). The catheter tip was advanced 8 cm caudally to the L1 vertebra and the other end of the catheter was sealed with a stainless-steel plug and affixed to the upper back. Animals were injected immediately with vehicle (PBS), mirin or KU-60019 followed by a 10μl PBS catheter flush. Injections were repeated every 24 h and drugs and vehicle reagents were delivered over 1 min time period using a Hamilton microlitre syringe (Hamilton Co, USA).

In a pilot experiment, mirin and KU-60019 were injected as described above at 1, 2, 5, 10 and 15 μg (n = 3 rats/group, 2 independent repeats) in a final volume of 10μl saline either daily, every other day or twice weekly for 28 d (not shown). Rats were then killed and the lesion site plus 5mm either side were harvested, lysed in ice cold lysis buffer, separated on 12% SDS PAGE gels and subjected to western blot detection of γH2Ax levels (Surey *et al.*, 2014). We determined that the amount of mirin and KU-60019 to optimally reduce γH2Ax levels by intrathecal delivery was 5 and 10 μg, respectively with a dosing frequency of every 24hrs. Optimal doses were then used for all experiments described in this manuscript. Rats were killed in a rising concentration of CO_2_ at either 28 d for immunohistochemistry and western blot analyses or 6 weeks for electrophysiology and functional tests.

### Knockdown of Mre11 using shRNA in vivo

SMARTvector Lentiviral shRNAs to Mre11 (shMre11; cat no. V3SR11242-240192989) and ATM (shATM; cat no. V3SR11242-238270626) under the control of a CMV promoter were purchased from Dharmacon and plasmid DNA was prepared from glycerol stocks according to the manufacturer’s instructions. A control plasmid containing the CMV promoter and a non-targeting control (shControl; cat no. VSC11721) was also purchased from Dharmacon. Plasmid DNA containing shMre11, shATM and shControl were complexed with *in vivo*-jetPEI (referred to as PEI from herein; Polyplus Transfection, New York, USA) according to the manufacturer’s instructions and injected into the DRG immediately after DC injury, as described by us previously (Jacques *et al.*, 2012; Almutiri *et al.*, 2018). We used *in vivo*-jetPEI, a non-viral vector, to deliver the shRNAs to DRGN *in vivo*, since we have previously shown that it transduces similar proportions of DRGN as adeno-associated virus 8 and that it does not require intra-DRG injection 1-2 week prior to injury to ensure maximum transgene expression (Almutiri *et al.*, 2018). In a preliminary experiment, we optimized the amount of plasmid DNA required to cause maximum suppression of γH2Ax^+^ foci in DRGN as 2μg for both shMre11 and shATM (not shown). Pre-optimised plasmid DNA was then injected intra-DRG and animals (n = 6/group/test, 3 independent repeats, total n = 18 rats/group/test) were killed after 28d for immunohistochemistry to detect γH2Ax^+^ or 6 weeks for electrophysiological and behavioural studies.

### Immunohistochemistry

Tissue preparation for cryostat sectioning and immunohistochemistry were performed as described previously (Surey *et al.*, 2014). Briefly, rats were intracardially perfused with 4% formaldehyde and optic nerves, eyes, L4/L5 DRG and segments of T8 cord containing the DC injury sites were dissected out and post-fixed for 2h at room temperature. Tissues were then cryoprotected in a sucrose gradient prior to mounting in optimal cutting temperature (OCT) embedding medium (ThermoFisher) and frozen on dry ice. Samples were then sectioned using a cryostat and immunohistochemistry was performed on sections from the middle of the optic nerve, DRG or spinal cord as described previously (Surey *et al.*, 2014). Sections were permeabilised in PBS containing 0.1 % Triton X-100, blocked in PBS + 3% w/v bovine serum albumin + 0.05% Tween-20 then stained with primary antibodies overnight at 4 °C. After washing in PBS, sections were incubated with secondary antibodies for 1 h at RT then washed further in PBS and mounted in Vectashield containing DAPI (Vector Laboratories). Primary antibodies used were: mouse anti-H2Ax pSer139 (γH2Ax; JBW301; 1:400 dilution; Merck) and rabbit anti-neurofilament (NF) 200 (1:400 dilution; Sigma); mouse anti-GAP43 (ThermoFisher; 1:400 dilution) was used to detect regenerating axons in the optic nerve and spinal cord. Regenerating axons in the DC were detected using GAP43 immunohistochemistry (Ahmed *et al.*, 2014; Almutiri *et al.*, 2018; Farrukh *et al.*, 2019) since Cholera toxin B labelling in our hands did not label regenerating axons by retrograde transport labelling in the rat (Ahmed *et al.*, 2014), despite others demonstrating successful labelling (Neumann and Woolf, 1999; Neumann *et al.*, 2002). Secondary antibodies used were Alexa-488 goat anti-mouse IgG and TexasRed goat anti-rabbit IgG (both from ThermoFisher). Controls were included in each run where the primary antibodies were omitted and these sections were used to set the background threshold prior to image capture. Image capture and analysis was performed by an investigator masked to the treatment conditions.

### Quantification of axon regeneration

Axon regeneration in the spinal cord was quantified according to previously published methods (Hata *et al.*, 2006). Briefly, serial parasagittal sections of cords were reconstructed by collecting all serial 50μm-thick sections (~70–80 sections/animal; n = 10 rats/treatment) and the number of intersections of GAP43^+^ fibers through a dorsoventral orientated line was counted from 6mm rostral to 4mm caudal to the lesion site. Axon number was calculated as a % of fibers seen 4mm above the lesion, where the DC was intact.

### Electrophysiology

Six weeks after surgery or treatment, compound action potentials (CAP) were recorded after vehicle, mirin and KU-60019 treatment as previously described (Lo *et al.*, 2003; Hains *et al.*, 2004; Almutiri *et al.*, 2018). Briefly, with the experimenter masked to the treatment conditions, silver wire electrodes were used apply single-current pulses (0.05ms) through a stimulus isolation unit in increments (0.2, 0.3, 0.6, 0.8, and 1.2 mA) at lumbar (L)1-L2 and compound action potentials (CAP) recorded at cervical (C)4-C5 along the surface of the midline spinal cord. CAP amplitudes were calculated between the negative deflection after the stimulus artefact and the next peak of the wave. CAP area was calculated by rectifying the negative component (full-wave rectification in Spike 2 software) and measuring its area at the different stimulation intensities. The dorsal half of the spinal cord was transected between stimulating and recording electrodes at the end of the experiment to confirm that a CAP could not be detected. Electrophysiology was analysed using Spike2 software (Cambridge Electronic Design, Cambridge, UK) and representative processed data are shown for CAP traces.

### Functional tests after DC injury

Functional testing after DC lesions was carried out as described by previously (Fagoe *et al.*, 2016; Almutiri *et al.*, 2018). Briefly, animals (n = 6 rats/group, 3 independent repeats (total n = 18/group/test)) randomly assigned and treatment status masked from the investigators, received training to master traversing the horizontal ladder for 1 week before functional testing. Baseline parameters for all functional tests were established 2-3 d before injury. Animals were then tested 2 d after DC lesion + treatment and then weekly for 6 weeks. Experiments were performed by 2 observers blinded to treatment with animals tested in the same order and at the same time of day. Three individual trials were performed each time for each animal.

#### Horizontal ladder crossing test

This tests the animals’ locomotor function and is performed on a 0.9-meter-long horizontal ladder with a diameter of 15.5 cm and randomly adjusted rungs with variable gaps of 3.5-5.0 cm. Animals were assessed traversing the ladder with the total number of steps taken to cross the ladder and the number of left and right rear paw slips being recorded. The mean error rate was then calculated by dividing the number of slips by the total number of steps taken.

#### Tape removal test (sensory function)

The tape removal test determines touch perception from the left hind paw. Animals were held with both hind-paws extended and the time it took for the animal to detect and remove a 15×15mm piece of tape (Kip Hochkrepp, Bocholt, Germany) was recorded and used to calculate the mean sensing time.

### Analysis of functional tests

The whole time-course of lesioned and sham-treated animals for the horizontal ladder crossing and mean tape sensing/removal test was compared using generalized linear mixed models (GLMM) or linear mixed models (LMM), as described previously (Fagoe *et al.*, 2016; Almutiri *et al.*, 2018). For the horizontal ladder test, we scored individual steps as either a successful step or a slip and therefore the data follows a binomial distribution. Data was compared using binomial GLMM, with lesioned/sham (‘LESION’; set to true in lesioned animals post-surgery, false otherwise) and operated/unoperated (‘OPERATED’; set to false before surgery, true after surgery) as fixed factors, animals as a random factors and time as a continuous covariate. Binomial GLMMs were then fitted in R using package *lme4* with the *glmer* function using the following model formulae (Fagoe *et al.*, 2016):

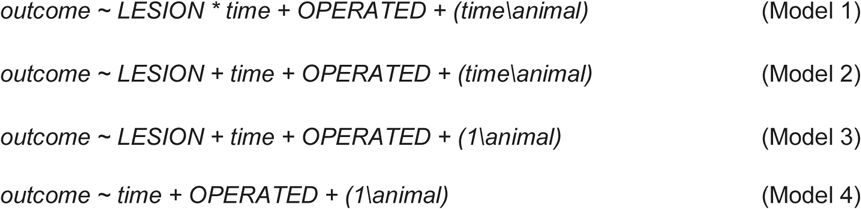

Bracketed terms refer to the ‘random effects’ and account for the presence of repeated measurements in estimation of the effect size and significance of INT and LESION. ‘Outcome’ for binomial GLMMs is a two-column list of counts of successes/fails per run as described in the individual models above. INT refers to the interaction term of LESION over time (Model 1). P-values for INT were then calculated, which represents differences in the evolution of outcomes over time, and LESION, the unconditional main effect of lesioning. Significance of specific parameters in LMMs and GLMMs can be determined by comparing a model containing the parameter of interest to a reduced model without it. Thus, INT was assessed by comparing Model 1 and Model 2, while LESION was assessed by comparing Model 3 and Model 4. LESION, represents the overall effect of lesioning on the outcome, while INT represents the difference in slope of the outcome between the two groups, i.e. speed of recovery. P-values for GLMMs were calculated by model comparison using parametric bootstrap for INT and LESION against the null hypothesis that each parameter is zero, using *pbkrtest* in R package. Between 1000 and 20000 simulations were used.

For tape removal test, the time-courses of lesioned versus sham were compared using linear mixed models (LMMs) with the R package *lme4* with the *glmer* function (Fagoe *et al.*, 2016; Almutiri *et al.*, 2018). Model formulae were the same as for the ladder crossing test above. Standard regression diagnostics (quantile plots of the residuals *vs* the normal distribution, plots of residuals *vs* fitted values) were carried out for the data fitted with LMMs. P-values for the INT and LESION parameters of the LMMs were calculated by model comparison using package *pbkrtest* in R, with the Kenward-Roger method. For the tape sensing and removal test, log of the withdrawal time was used as the data were expected to follow an exponential distribution. Independent sample T-tests were performed to determine statistical differences at individual time points.

### Analysis of pH2A positive foci

#### Drosophila brains

Adult fly brains were dissected in PBS and fixed and stained as described above. 41 × 41 × 13.5μm volumes of the same region of the central brain were imaged at 16-bit depth for each brain using a 63x water immersion N.A. 1.2 objective on a Zeiss LSM880 confocal microscope. The zoom was set at 3.2 and z-step at 0.45 μm and Airy scan processing module used with super-resolution settings to improve spatial resolution post-collection. To display visualise brightly-staining pH2Av^+^ foci only, the lower intensity pan-nuclear staining was eliminated by manually resetting the black level threshold in Fiji-3. The identical original and processed images are displayed together in supplementary figures. Movies were rendered in Fiji-3.

#### DRGN

1. Frequency distribution: Images of the entire DRG in the middle 3 sections (n = 160 images/DRG) of L4/L5 DRG from each animal (n = 10) were captured by an investigator masked to the treatment conditions at x10 magnification using a Zeiss Axioplan 200 epifluorescent microscope. Images were merged in Adobe Photoshop using Photomerge and the frequency of γH2Ax^+^ foci in different diameter DRGNs was recorded (Jacques *et al.*, 2012).
2. Relative fluorescent staining intensity: the integrated density of fluorescence was measured using ImageJ as previously described (Surey *et al.*, 2014). Briefly, images from the middle 3 DRG sections from n = 10 rats were thresholded and the mean integrated density of pixels/cell was recorded.
3. Western blot: Total protein was extracted from pooled L4/L5 DRG pairs from n = 3 rats/group (i.e. 6 DRG) after DC+vehicle, mirin and KU-60019 treatment and western blots followed by subsequent densitometry was performed as described by us previously and in brief below (Ahmed *et al.*, 2005). Experiments were repeated on 3 independent occasions (total n = 9 rats/group i.e. 18 DRG/group).

### Western blots

Proteins were extracted from hippocampal, retinal or DRG neuron cultures/tissues as described by us previously (Ahmed *et al.*, 2005). Briefly, cultures were treated with ice-cold lysis buffer (20mM HEPES pH 7.5, 1 mM EDTA, 150 mM NaCl, 1% NP-40 and 1 mM dithiothreitol supplemented with protease (Roche, Welwyn Garden City, UK) and phosphatase (ThermoFisher) inhibitor cocktails), protein concentration was determined by Bradford Protein Assay and 15 μg of total protein separated on 12% Tris-glycine SDS-PAGE gels. Proteins were transferred to nitrocellulose membranes, blocked with 2.5% non-fat dry milk in TBS containing 0.05% Tween-20 for 90 min with agitation then incubated for 2 h with primary antibodies and for 1 h with secondary antibodies diluted in 2.5% non-fat dry milk at RT. For hippocampal neurons, proteins were detected using the Odyssey imaging scanner (LI-COR). Primary antibodies used were: mouse anti-Synapsin antibody (Santa Cruz) and mouse anti-actin antibody (Sigma, A5316). Secondary antibodies used were: Alexa-680 goat anti-mouse IgG and Alexa-800 goat anti-rabbit IgG (New England Biolabs). For DRGN cultures, membranes were probed with primary anti-γH2Ax (pSer139 monoclonal antibody; 1:400 dilution; Merck) and HRP-labelled anti-mouse secondary antibody before bands being detected using an enhanced chemiluminescence kit (GE Healthcare, Buckingham, UK). β-actin (Sigma: 1:1000 dilution) was used as a protein loading control for western blots.

### Densitometry

Western blots were scanned into Adobe Photoshop (Adobe Systems, San Jose, CA, USA) keeping all scanning parameters constant between blots. The integrated density of bands was analysed using the built-in macros for gel analysis in ImageJ as described by us previously (Ahmed *et al.*, 2006a).

### Statistical analysis

All data are presented as means ± standard error of the mean (SEM). Comparison of means were performed using one-way or two-way ANOVA using Prism 7 (San Diego, CA, USA) or SPSS (Version 25, IBM, New York, USA). Tukey’s significant difference test followed ANOVA’s, as appropriate. In Fig. 2A a Kruskal-Wallis test with Dunn’s post-hoc was performed. Functional tests in rats were analysed using R package (www.r-project.org) as described above. Tests used for each experiment are described in the corresponding methods. In all tests, *P*<0.05 were considered statistically significant. Supplementary Table 1 lists all tests performed, the comparisons made, and the associated *P*-values.

## Data availability

The authors confirm that the data supporting the findings of this study are available within the article and its Supplementary material.

## Results

Genetic targeting of the double strand break-sensing MRN complex is neuroprotective in *Drosophila.* Double strand breaks are a feature of early-stage Alzheimer’s disease and correlate with reduced cognitive score (Simpson *et al.*, 2015). Double strand breaks are also generated in neurons *in vivo* exposed to Aβ_1-42_ oligomers (Suberbielle *et al.*, 2013). Hence, we chose to target the MRN complex in a *Drosophila* model of Alzheimer’s model as proof-of-concept. In this model, tandem Aβ_1-42_ oligomers for Alzheimer’s disease (AD) were expressed in, and secreted from, adult post-mitotic neurons to replicate the extracellular deposition in AD (Speretta *et al.*, 2012). Expression was restricted to adult neurons only to rule out confounding developmental phenotypes (see Methods for genetics). First, we confirmed that double strand breaks were generated in neurons exposed to A oligomers, as predicted from other studies and our own studies in rat hippocampal neurons (Fig. 2). Activation and recruitment of ATM to the site of double strand breaks lead to multiple phosphorylation events, one of which is the phosphorylation of histone γH2Ax at and around the site of the break, resulting in the formation of characteristic γH2Ax^+^ nuclear foci (Shrivastav *et al.*, 2008). Therefore, antibodies to γH2Ax (H2Av in *Drosophila*) are widely use to visualise the histone modifications surrounding a double strand break (Celeste *et al.*, 2002; Simpson *et al.*, 2015; Milanese *et al.*, 2018). Anti-pH2Av staining of control brains revealed neurons with a single brightly staining focus per nucleus corresponding to the nucleolus organizing centre. In contrast, many of the Aβ_1-42_-expressing neurons had multiple, clustered foci (Fig. 1A, Supplementary Fig. 1 and Movie 1).

**Figure 1.**
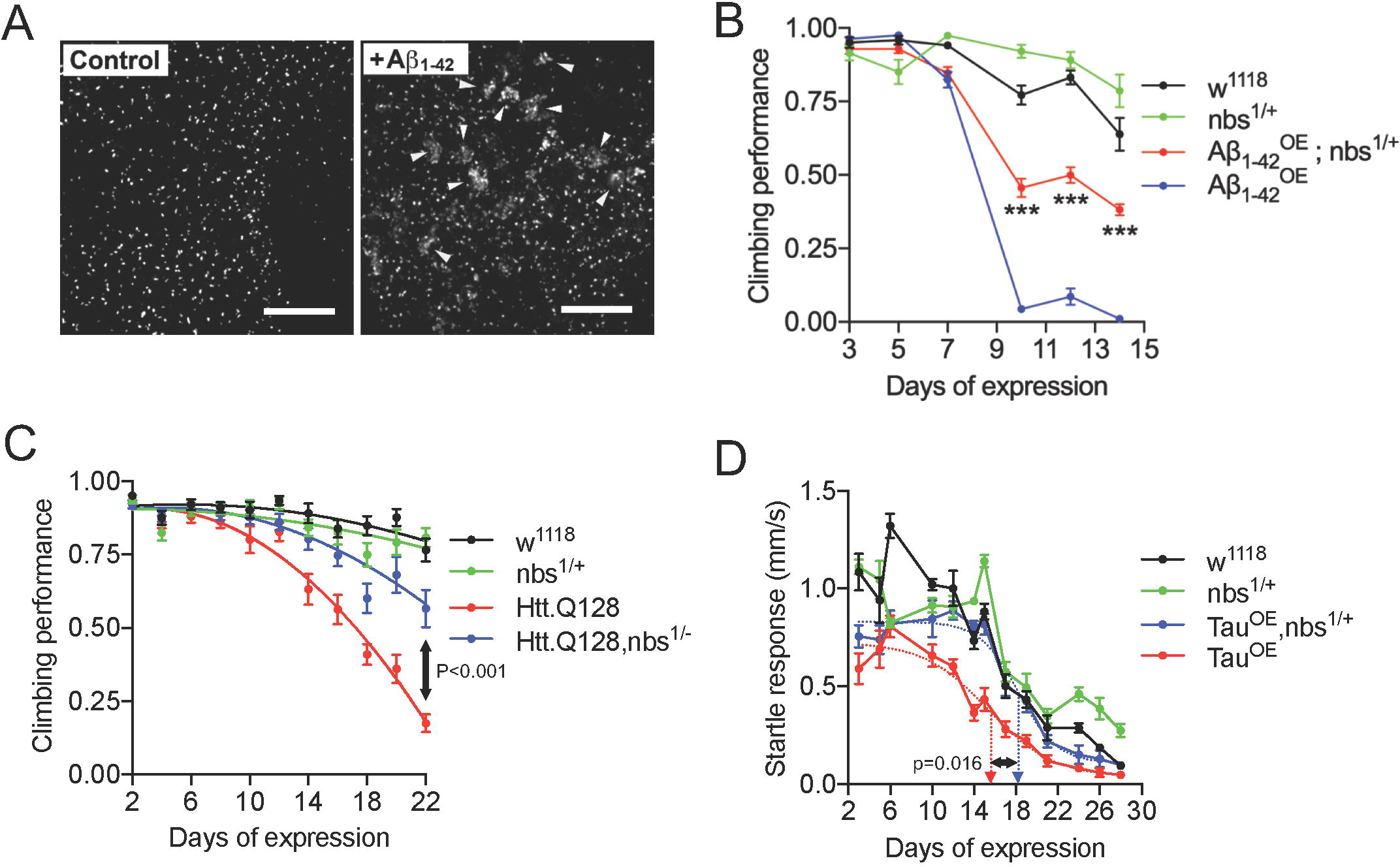
Targeting the MRN complex is neuroprotective in *Drosophila* models of neurodegeneration. (**A**) Expression of tAβ_1-42_ in adult neurons generates DNA double-strand breaks. Projections of central brain neurons stained with anti-pH2Av. Control neurons display a single pH2Av^+^ focus per nucleus corresponding to the nucleolus organising region. Multiple neurons expressing Aβ_1-42_ display large numbers of clustered foci (arrowheads). (**B**) Flies were startled by tapping to the base of a vial and the negative geotaxis climbing response quantified as a measure of neural output. Flies expressing (**B**) Aβ_1-42_ or (**C**) Htt.Q128 in adult neurons show a rapid decline in climbing ability which is partially suppressed in flies heterozygous for a null allele of *nbs*. Statistical comparisons in (**B**) by 2-way ANOVA and in (**B**) by extra sum-of-squares F-test, *** indicates p<0.001. (**D**) When stimulated by vibration, Tau-expressing flies show a rapid decline in the startle response which is partially suppressed in *nbs^−/+^* flies. Dashed arrows indicate days of expression required for 50% decline in response determined from fitted lines; comparison by 2-tailed t-test. Scale bars in (**A**) = 10µm.

**Figure 2.**
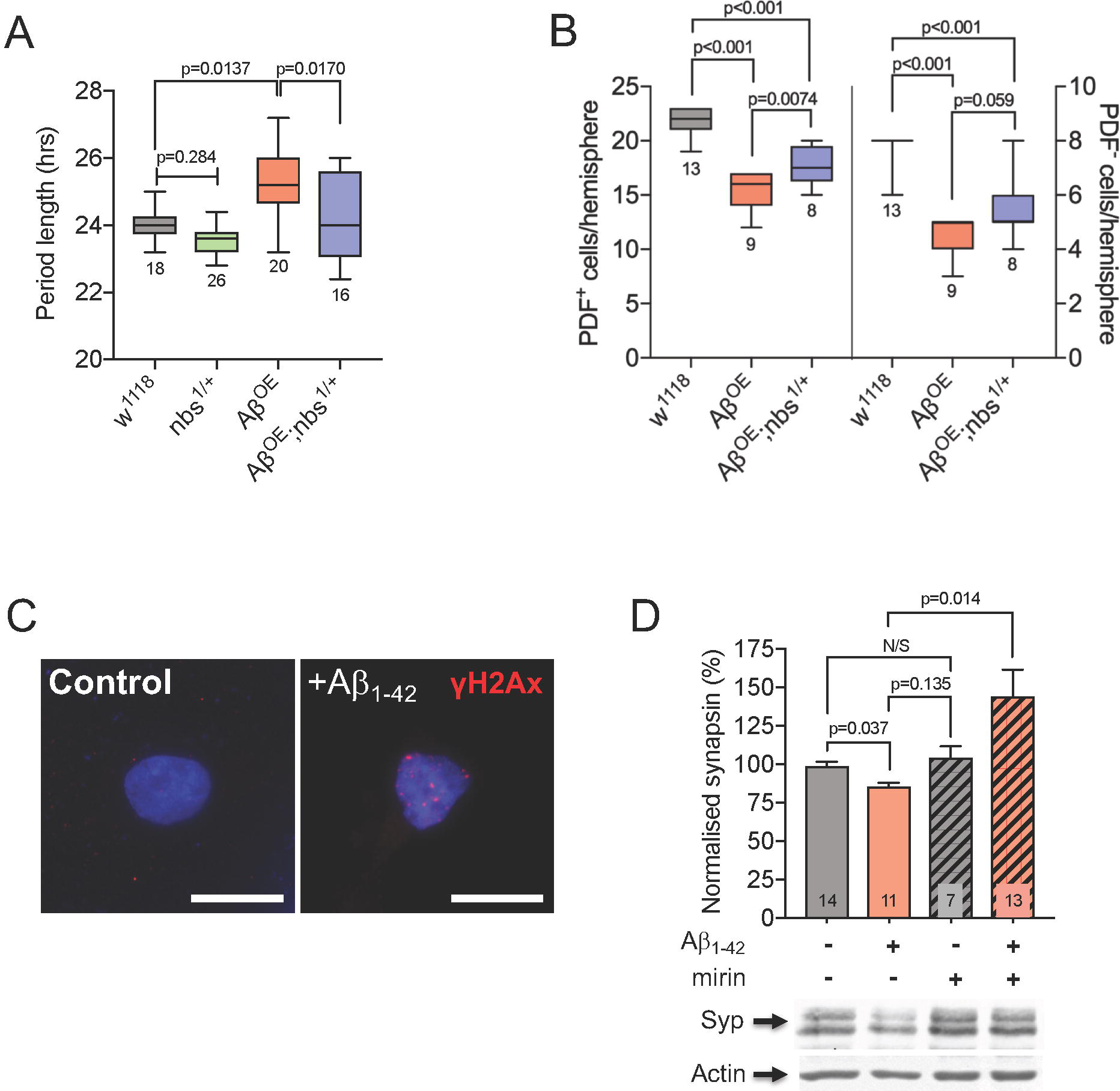
Targeting the MRN complex protects against Alzheimer’s disease-relevant phenotypes. (**A**) Expression of Aβ_1-42_ in clock neurons leads to loss of Per^+^ cells. In *nbs^−/+^* flies, loss of cells is partially prevented (Kruskal-Wallis with Dunn’s post-hoc test; p values and n are indicated). (**B**) The free-running circadian locomotor cycle of control flies is ∼24 h but is considerably lengthened when Aβ_1-42_ is expressed in clock neurons indicating weaker rhythmicity in the circadian circuitry. The increase is suppressed in *nbs^−/+^* flies (CLEAN spectral analysis; comparisons by ANOVA with Tukey’s post-hoc test; p values and n are indicated). (**C**) Exposure of rat hippocampal neurons to Aβ_1-42_ oligomers *in vitro* generate double strand breaks that can be visualized by staining with anti-γH2Ax (red). DNA is visualized with DAPI (blue). (**D**) Quantification of synapsin levels in hippocampal neurons by Western blot. Exposure to Aβ_1-42_ oligomers leads to loss of the pre-synaptic protein, synapsin, which is reversed by the Mre11 inhibitor, mirin (mean ± SEM; ANOVA with Tukey’s post-hoc test; P values and n are indicated). Scale bars in (**C**) = 10µm.

Next, we asked whether reducing *nbs* levels genetically would be neuroprotective in Aβ_1-42_ expressing flies. We chose to introduce one null allele of *nbs* to reduce the gene dosage by 50% and then used the ability of flies to climb after being startled as a measure of neural output. This negative geotaxis assay is widely used in *Drosophila* neurodegeneration studies: climbing ability declines rapidly and correlates with a reduced speed of neural transmission (Kerr *et al.*, 2011). In flies expressing Aβ_1-42_ oligomers, climbing ability decreased rapidly with age yet this was partially suppressed in *nbs^−/+^* heterozygous flies (Fig. 1B and see Movie 2). The toxicity of the less aggregative tAβ_1-42_ construct lacking the 12 amino acid flexible linker was similar suppressed in *nbs^1/+^* heterozygous flies and a second null allele, nbs^2/+^ showed a similar effect (Supplementary Fig. 2A and B).

To confirm that the neuroprotective effect was not specific to Aβ_1-42_ pathology, we used the same approach for flies expressing an expanded Htt protein associated with Huntington’s disease (Htt.Q128) (Romero *et al.*, 2008) and saw a similar neuroprotective effect in climbing assays (Fig. 1C). Again, to confirm the suppression was not specific to climbing ability, we switched to a tracking system capable of monitoring the horizontal movement of flies continuously and expressed the human 0N4R Tau carrying the R406W point mutation associated with frontal temporal dementia with Parkinsonism (Wittmann *et al.*, 2001). Tau-expressing flies moved at normal speed when unstimulated (not shown) but their escape response declined rapidly with age. Again, this was partially suppressed in *nbs^−/+^* flies (Fig. 1D) and this neuroprotective effect was not due to a reduction in Tau levels nor to changes in Tau phosphorylation in the brains of these flies (Supplementary Fig. 2C). Finally, to ensure these protective effects were not specific to *nbs*, we depleted *rad50* in Htt.Q128-expressing flies using the same strategy of reducing the gene dosage with a null allele. Flies with reduced *rad50* (*rad50^−/+^*) partially suppressed the decline in climbing ability in the Htt.Q128 flies (Supplementary Fig. 2D). We also expressed expanded HttQ128 in the developing eye under the control of *GMR-gal4*, which generates a progressive degeneration evidenced as a widespread loss of pigmentation after 6-7 weeks (Romero *et al.*, 2008). In *rad50^−/+^* flies, pigmentation is restored, indicating enhanced cellular survival (Supplementary Fig. 2E).

### Targeting the MRN complex suppresses neurodegeneration-relevant phenotypes

AD patients commonly suffer disrupted circadian behaviour patterns (Saeed and Abbott, 2017) and a similar effect is observed in Aβ-expressing flies (Long *et al.*, 2014). We asked whether reducing *nbs* genetically would suppress this more directly disease-relevant phenotype. Tandem Aβ_1-42_ was expressed specifically in clock neurons under the control of *tim-Gal4* and their survival studied with immunocytochemistry (Renn *et al.*, 1999). Aβ-expression induced loss of some Per^+^ neurons but this was partially suppressed in *nbs^−/+^* flies (Fig. 2A). Consistent with the loss of clock neurons (Renn *et al.*, 1999), Aβ-expressing flies exhibited significantly longer free-running circadian locomotor cycles than controls but the normal ∼24 h periodicity was restored in *nbs^−/+^* flies (Fig. 2B and Supplementary Fig. 2F).

We extended our findings to mammalian neurons by examining synapse loss – a common early feature of neurodegenerative diseases (Gillingwater and Wishart, 2013). Cultured primary hippocampal neurons were exposed to Aβ_1-42_ oligomers and stained with anti-γH2Ax antibodies. As expected, the characteristic foci corresponding to double strand breaks in the nuclei were induced by exposure to Aβ_1-42_ (Fig. 2C). Synapsin levels were significantly reduced 24 h after addition of Aβ oligomers (Fig. 2D) suggesting that one route to synapse loss in AD might be via Aβ-induced double strand breaks triggering the DNA damage response. Mirin, a small-molecule Mre11 exonuclease inhibitor (Dupre *et al.*, 2008), prevented synapsin loss (Fig. 2D), indicating that targeting the MRN complex can also protect mammalian neurons from changes associated with early-stage neurodegeneration.

### Mre11 or ATM inhibition prevents apoptosis and stimulates regeneration after neurotrauma

Double strand breaks are a feature of acute neurological disorders (Hayashi *et al.*, 1998; Kotipatruni *et al.*, 2011). DNA strand breaks have been observed in retinal ganglion cells (RGC) after optic nerve injury (He *et al.*, 2012) and neurons suffering from ischemia display genome fragility and fragmented DNA both *in vitro* and *in vivo* (Yang *et al.*, 2016). We used an adult rat primary retinal culture system enriched in RGC, where cells are grown in the presence of inhibitory CNS myelin extracts to simulate a post-injury environment (Ahmed *et al.*, 2006b). As expected, we detected the presence of double strand breaks with anti-γH2Ax antibodies in the RGC (Fig. 3A and B). The RGC die rapidly by apoptosis and, even if apoptosis is blocked with inhibitors, they fail to regenerate neurites (Vigneswara *et al.*, 2013). We treated the RGC with mirin to attenuate the DNA damage response. This blocked RGC apoptosis and dramatically stimulated neurite regrowth. Mirin treatment was significantly more effective than a current positive control treatment: ciliary neurotrophic factor (CNTF) (Fig. 3C and D). Since a key function of the MRN complex is to recruit and activate ATM at double strand breaks (Lee and Paull, 2005), and given that HD-pathology is reduced by targeting ATM (Lu *et al.*, 2014), we asked whether inhibiting ATM directly would also be neuroprotective to RGC. Treatment with the highly selective ATM inhibitor, KU-60019, had similarly dramatic effects on RGC survival and neurite outgrowth as mirin (Fig. 3C-F).

**Figure 3.**
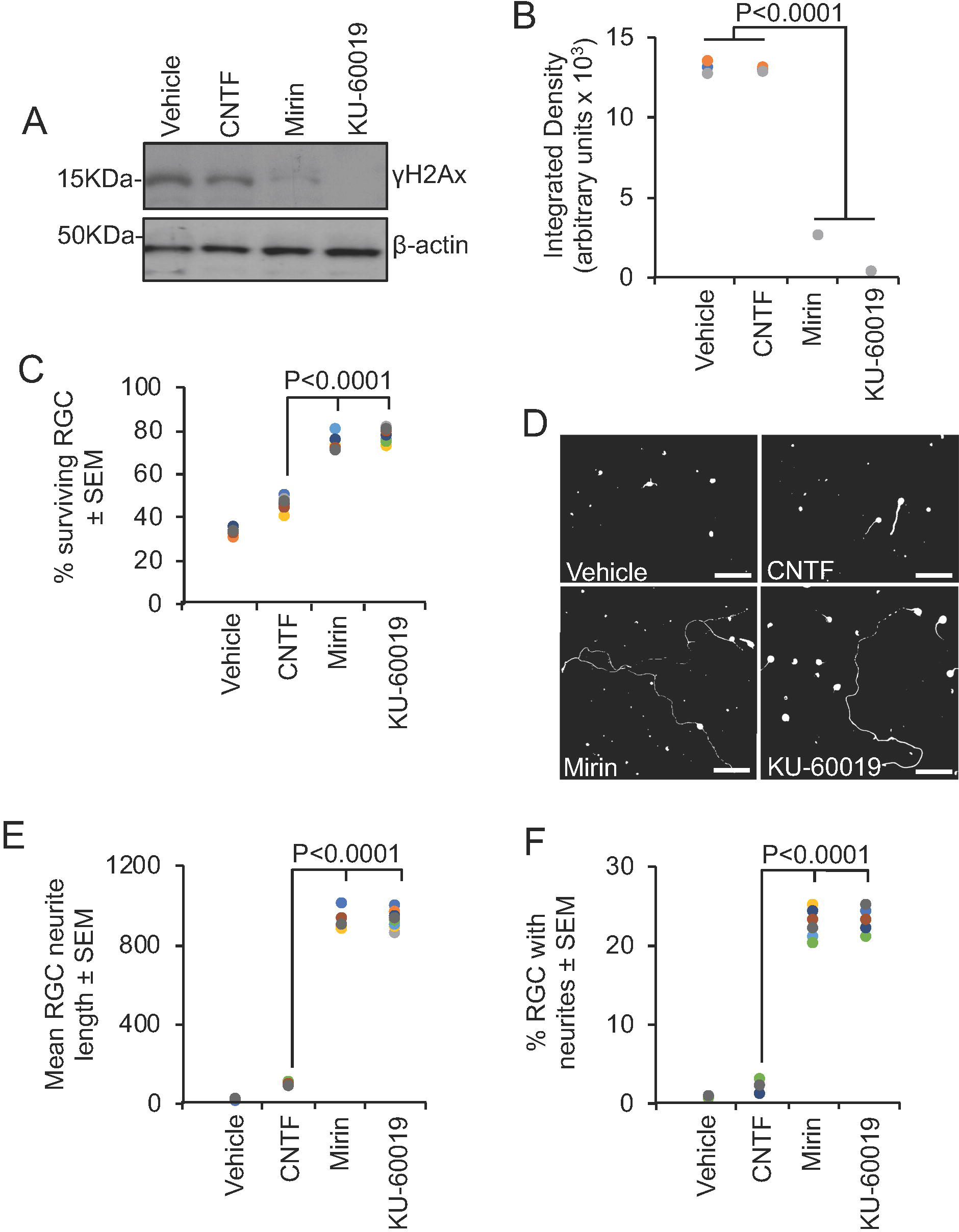
Inhibition of Mre11 prevents RGC apoptosis and stimulates neurite outgrowth after 4 days in culture in the presence of inhibitory CNS myelin extracts. (**A**) Western blot and (**B**) subsequent densitometry to show attenuated levels of anti-γH2Ax after treatment with mirin and KU-60019. (**C**) Mirin or KU-60019 significantly enhanced RGC survival. (**D**) Representative images from RGC treated with vehicle, CNTF (positive control), mirin and KU-60019. Mirin or KU-60019 treatment (**E**) increased the mean RGC neurite length and (**F**) % RGC with neurites. n = 3 wells/treatment, 3 independent repeats (total n = 9 wells/condition). AU = arbitrary units. Comparisons in (**B**), (**C**), (**E**) and (**F**) by one-way ANOVA with Dunnett’s post-hoc test. Scale bars in (**D**) = 100µm.

We extended our *in vitro* findings to an *in vivo* optic nerve crush injury paradigm. We observed low levels of γH2Ax in intact control rat retinae *in vivo* (Fig. 4A and B). However, high levels of γH2Ax were observed within 1 d and at 24 d after optic nerve crush injury, suggesting rapid activation of double strand breaks and persistence of high levels for the duration of the experiment i.e. 24 days. Thus, the DNA damage response pathway was persistently activated in the retina post-injury. We used a pre-optimized dosing regimen for mirin and KU-60019 for 24 days after optic nerve crush injury to attenuate the DNA damage response, as demonstrated by significantly suppressed γH2Ax in western blots (Fig. 4C) and subsequent densitometry (Fig. 4D) and saw unprecedented RGC survival of 93% and 91%, respectively, when compared to intact controls (Fig. 4E and F). Both mirin and KU-60019 also promoted significant numbers of RGC to regenerate their axons, which emerged from the lesion site and grew for long distances (Fig. 4G and H). The numbers of regenerating axons we see are, to the best of our knowledge, unprecedented when compared to known treatments and the axons also regenerate longer distances into the distal optic nerve than has been seen previously (Berry *et al.*, 1996; Leon *et al.*, 2000; Vigneswara *et al.*, 2013).

**Figure 4.**
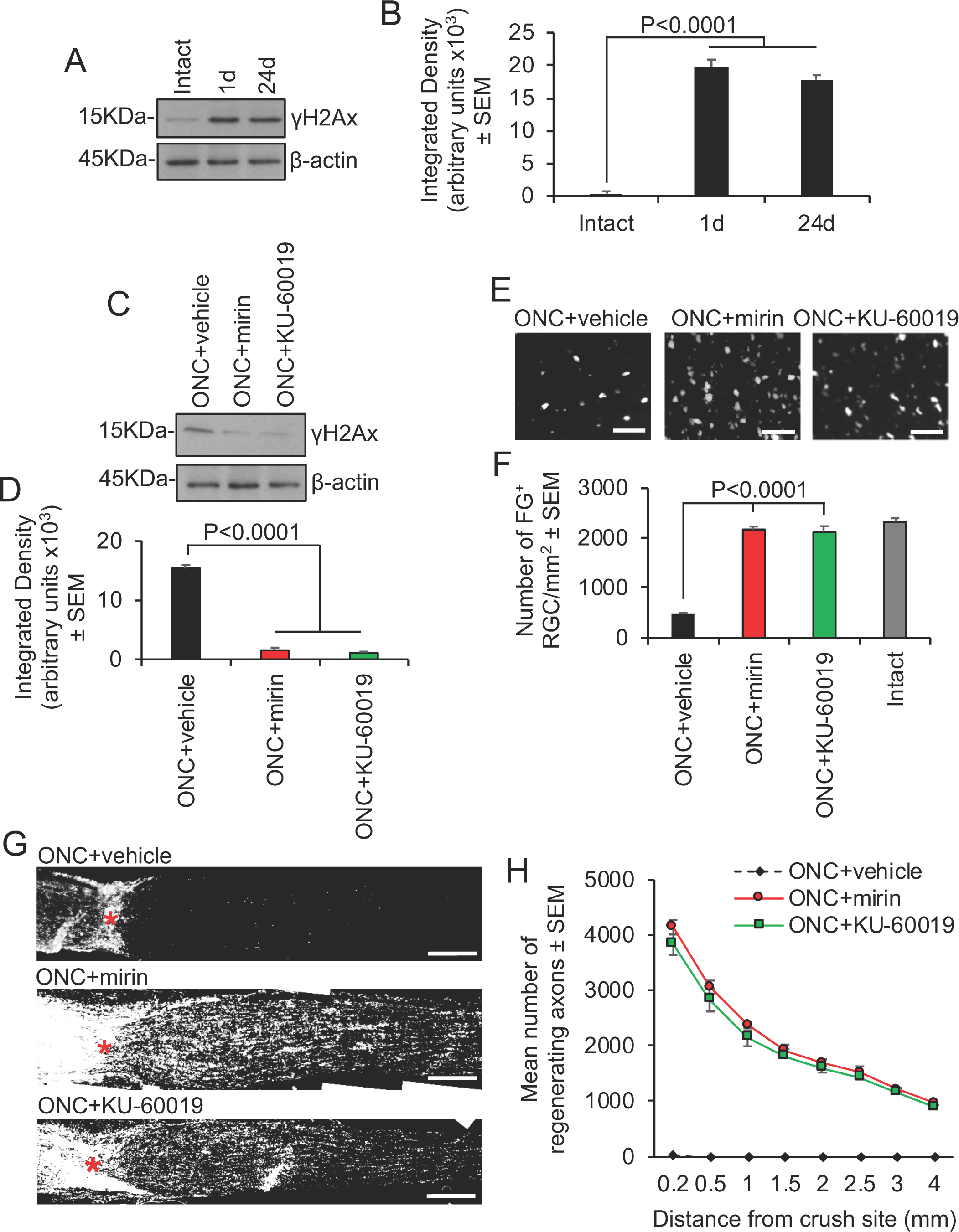
Inhibition of Mre11 and ATM prevents RGC apoptosis and stimulates axon regeneration at 24 days after optic nerve crush injury. (**A**) Western blot and (**B**) subsequent densitometry to show phosphorylation of H2Ax (γH2Ax) as a marker of DNA damage after optic nerve crush injury (n = 6 retinae/time-point, 3 independent repeats (total n = 18 retinae/time-point)). (**C**) Western blot and (**D**) densitometry to show that mirin and KU-60019 significantly suppress optic nerve crush injury-induced γH2Ax levels (n = 6 retinae/time-point, 3 independent repeats (total n = 18 retinae/condition). (**E**) Representative images and (**F**) quantification of FluoroGold backfilled RGC in retinal wholemounts to demonstrate that mirin and KU-60019 significantly enhanced RGC survival at 24 days after optic nerve injury (n = 6 retinae/time-point, 3 independent repeats (total n = 18 retinae/condition). (**G**) representative images and (**H**), quantification to show that mirin and KU-60019 significantly enhanced RGC axon regeneration as detected by GAP43 immunoreactivity (n = 6 nerves/condition, 3 independent repeats (total n = 18 nerves/condition). AU = arbitrary units. Comparisons in (**B**), (**D**), (**F**) and (**H**) by one-way ANOVA with Dunnett’s *post-hoc* test. Scale bars in (**E**) = 50µm and in (**G**) = 200µm).

### Mre11 and ATM inhibitors promote DRGN survival and neurite outgrowth *in vitro* and DC axon regeneration after spinal cord injury *in vivo*

Double strand breaks are also generated in spinal neurons after spinal cord injury (SCI) (Kotipatruni *et al.*, 2011) and there is an urgent need for effective therapies to treat SCI patients. Initially, we tested our DNA damage response attenuation strategy using a similar *in vitro* culture model to the retina in which dorsal root ganglion neurons (DRGN) containing double strand breaks are cultured from adult rats and grown in the presence of inhibitory CME (Supplementary Fig. 3A). Again, mirin and KU-60019 treatment stimulated significant DRGN survival and promoted neurite outgrowth (Supplementary Fig. 3B-E).

To generate a spinal cord injury *in vivo*, we surgically injured the ascending long tract axons of the dorsal funiculus in adult rats by dorsal column (DC) crush injury (Surey *et al*., 2014) and mirin or KU-60019 inhibitors were administered through an intrathecal catheter. None of the animals showed adverse effects to SCI and all animals were included for analysis. Double strand breaks formed in DRGN of all diameters after DC+vehicle treatment, as predicted from previous studies (Kotipatruni *et al.*, 2011) (Fig. 5A and B). However, quantification of the intensity of γH2Ax^+^ was significantly reduced in DC+mirin and DC+KU-60019-treated animals (Fig. 5C). Similarly, western blot and subsequent quantification for γH2Ax levels showed significant attenuation in DC+mirin and DC+KU-60019-treated rats, indicating partial suppression of the DNA damage response (Fig. 5D and E).

**Figure 5.**
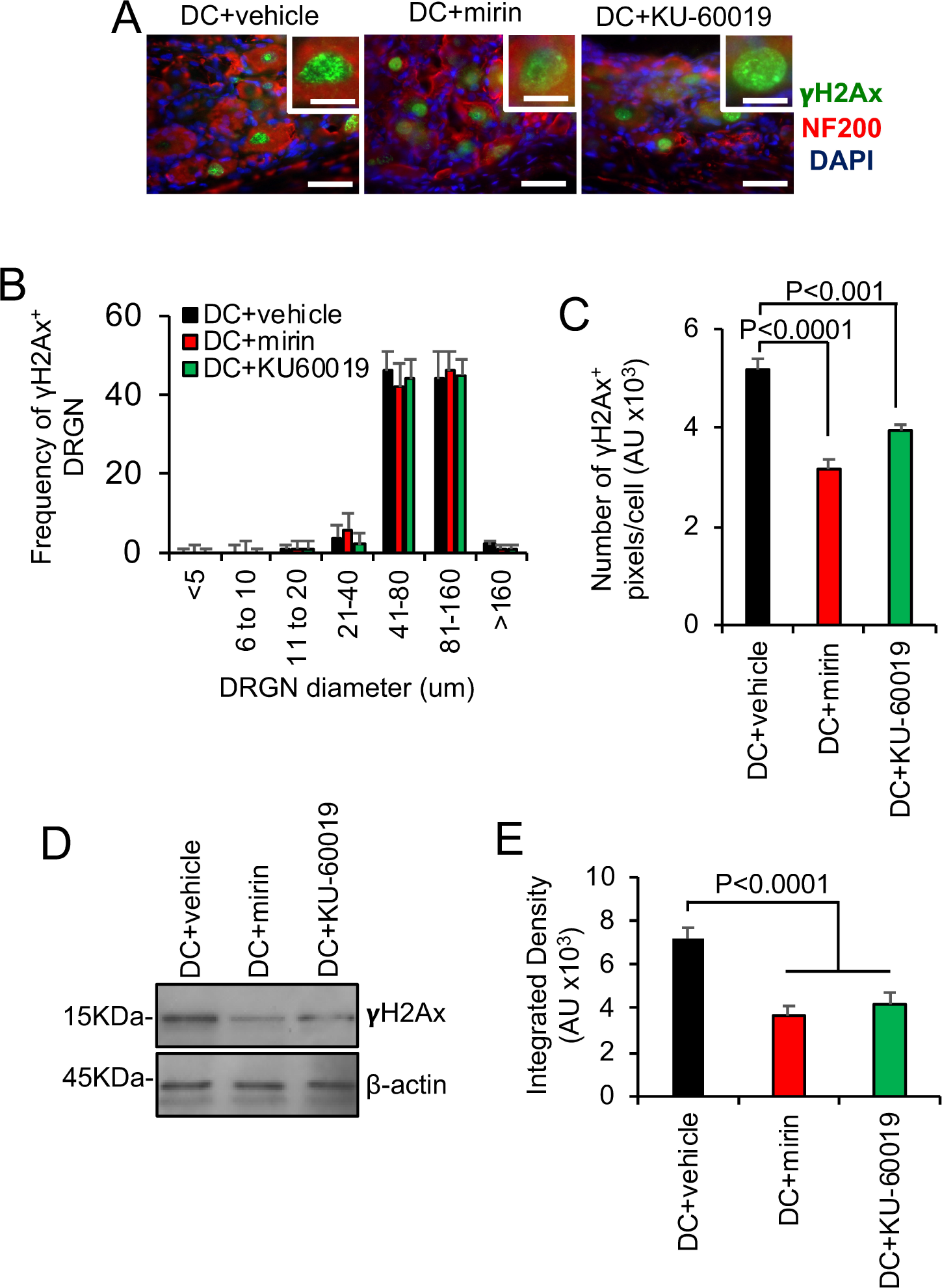
Inhibition of Mre11 and ATM reduces the number of DRGN with double strand breaks and the levels of γH2Ax after DC injury *in vivo*. (**A**) Representative images from DC+vehicle, DC+mirin and DC+KU-60019-treated sections of rat DRGN (n = 6 rats/group, 3 independent repeats; total n = 18 rats/group) demonstrating γH2Ax (green) localization in the nucleus of DRGN (red). Images are counterstained with DAPI (blue) to demarcate the nucleus. (**B**) Quantification of the frequency of γH2Ax^+^ DRGN sorted by different soma size. (**C**) Quantification of the mean pixel intensity/DRGN in DC+vehicle, DC+mirin and DC+KU-60019-treated DRGN. (**D**) Western blot and (**E**) subsequent densitometry to demonstrate reduction of γH2Ax protein levels in DRGN after treatment with mirin and ATM inhibitors. AU = arbitrary units. Comparisons in (**C**) and (**E**) by one-way ANOVA with Dunnett’s post-hoc test. n = 6 rats/treatment, 3 independent repeats (total n = 18 rats/treatment). Scale bars in **A** = 50µm, insets in **A** = 10µm.

We next investigated if suppression of DNA damage response by mirin and KU-60019 promotes DC axon regeneration after injury in rats. Little or no GAP43^+^ immunoreactivity was detected in DC+vehicle-treated rats whilst a large cavity was present at the lesion site (#) (Fig. 6A). Quantification of the % of axons in DC+vehicle-treated rats falls rapidly to 0% at the lesion site (i.e. 0mm) and remains at 0% for all distances quantified (Fig. 6B). In DC+mirin and DC+KU-60019-treated rats, significant GAP43^+^ immunoreactivity was present beyond the lesion cavity and in the rostral segment of the spinal cord (Fig. 6A and B). Quantification of the % of axons demonstrated significantly enhanced numbers at all distances when compared to DC+vehicle-treated rats, with DC+KU-60019 treatment regenerating slightly greater numbers of GAP43^+^ axons at all distances rostral to the lesion (Fig. 6B). For example, DC+KU-60019 treatment regenerated 47 ± 8, 35 ± 6, 25 ± 4 and 18 ± 3% of axons at 0, 2, 4 and 6mm rostral to the lesion site, compared to DC+vehicle-treated rats (Fig. 6B).

**Figure 6.**
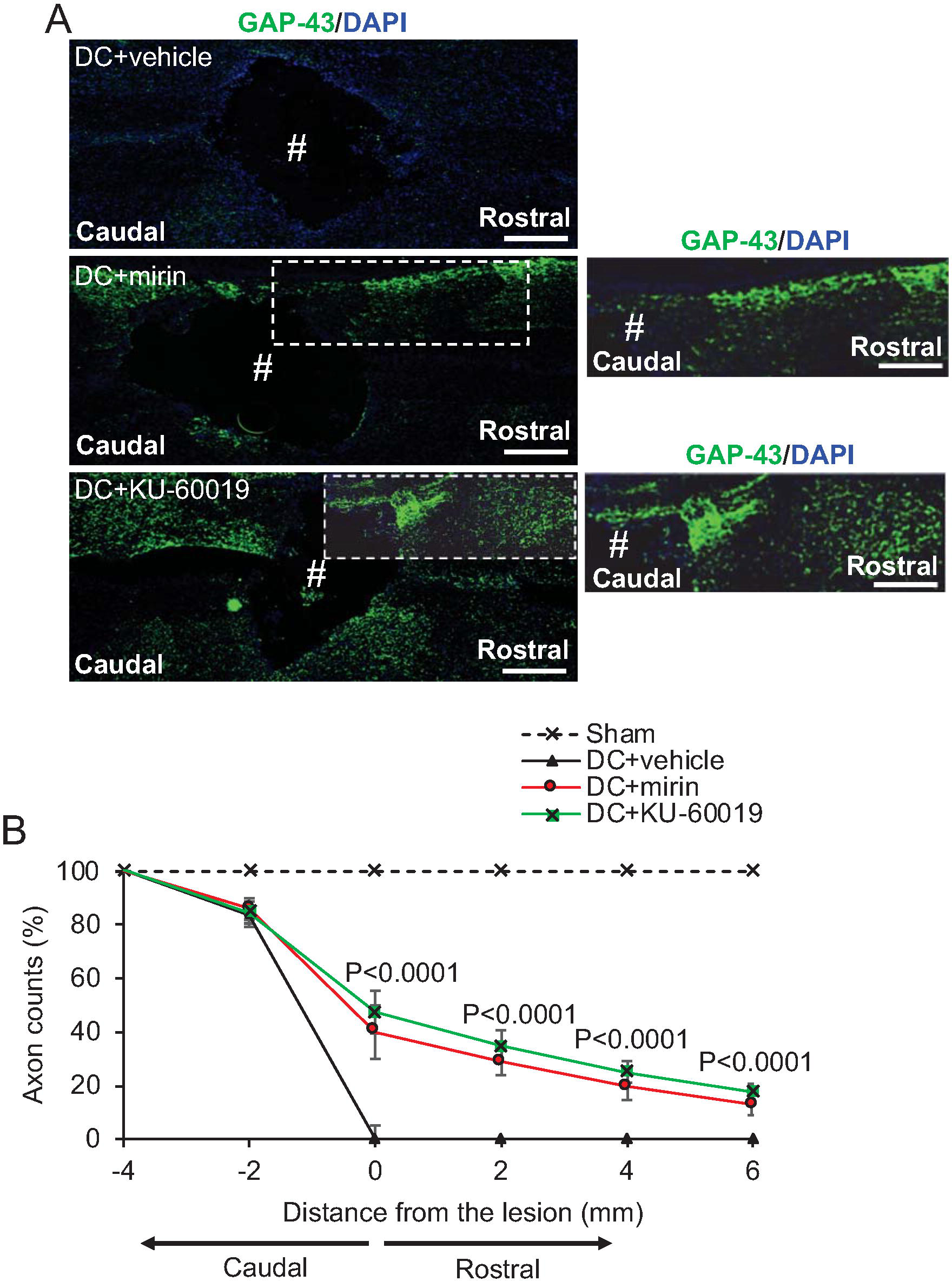
Intrathecal injection of Mre11 and ATM inhibitors promote significant axon regeneration at 6 weeks after spinal cord injury. (A) GAP43^+^ (green) immunoreactivity was largely absent in DC+vehicle-treated groups (Blue = DAPI^+^ nuclei). However, in DC+mirin and DC+KU-60019-treated rats, enhanced GAP43^+^ immunoreactivity was present in the rostral segment of the spinal cord in beyond the lesion site (#) where a large cavity remained. (**B**) Quantification of the % of axons at different distances rostral to the lesion site showed significantly enhanced numbers of GAP43^+^ regenerating axons at all distances tested after treatment with DC+mirin and DC+KU-60019. (n = 10 nerves/condition). Scale bars in (**A**) = 200µm. Comparisons in (**B**) by one-way ANOVA with Dunnet’s *post-hoc* test (DC+vehicle *vs* DC+KU-60019).

### Mre11 and ATM inhibitors restore function after spinal cord injury *in vivo*

We employed electrophysiology and simple functional tests (Almutiri *et al.*, 2018) to quantify recovery from the injury after mirin or KU-60019 treatment. In DC+vehicle-treated rats, the normal compound action potential (CAP) trace across the lesion site in Sham controls was abolished, whilst treatment with either mirin and KU-60019 restored 50-55% of the CAP trace (Fig. 7A) and the CAP amplitude (Fig. 7B) observed in sham-treated animals. As expected, dorsal hemisection between the stimulating and the recording electrodes at the end of the experiment abolished CAP traces in all animals (if a CAP trace was observed), confirming that the experiment was technically successful (Fig. 7A). The CAP area was also significantly improved in mirin and KU60019-treated rats compared to vehicle groups (Fig. 7C). The improved electrophysiological function translated into very dramatic improvements in sensory and locomotor function. We used a tape sensing-and-removal test and a horizontal ladder-crossing test to quantify recovery of sensory and motor function, respectively (Almutiri *et al.*, 2018). In sham-treated animals, there is a small initial decline in mean sensing/removal time (Fig. 7D) and mean ratio of slips to total steps (Fig. 7E) due to the effects of anaesthetic but these are restored to baseline levels within 3 weeks. Vehicle-treated groups displayed significantly impaired sensory and motor function that failed to return to baseline levels even after 6 weeks. In contrast, mirin or KU-60019-treated groups both showed dramatically improved sensing times and ladder-crossing ability 2d after surgery and after 3 weeks were indistinguishable from the sham-treated control. These results demonstrate that either Mre11 or ATM inhibition is able to promote functional recovery after SCI *in vivo*, which we believe to be at unprecedented levels.

**Figure 7.**
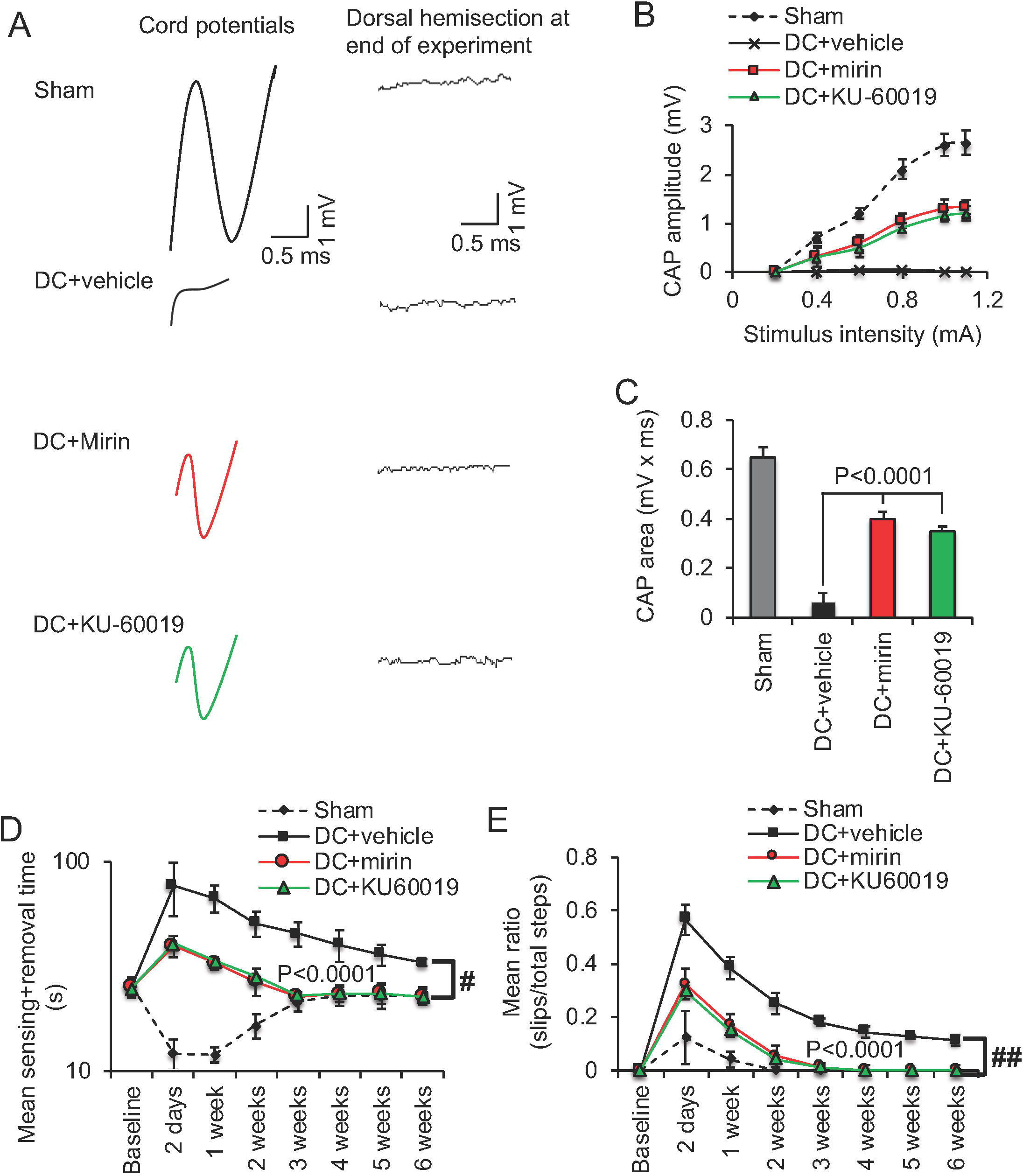
Intrathecal injection of Mre11 and ATM inhibitors promotes significant functional repair after spinal cord injury. (**A**) Spike 2 software processed CAP traces from representative sham controls, DC+vehicle, DC+mirin and DC+KU-60019-treated rats showing some recovery of dorsum cord potentials in mirin and KU-60019-treated rats. Dorsal hemisection between the stimulating and the recording electrodes at the end of the experiment ablated CAP traces in all animals demonstrating technical success of the experiment. (**B**) Negative CAP amplitudes were significantly attenuated in DC+vehicle-treated rats but were significantly improved in DC+mirin and DC+KU-60019-treated rats (P<0.0001, one-way ANOVA (main effect)). (**C**) Mean CAP area at different stimulation intensities were significantly attenuated in DC+vehicle-treated rats but improved significantly in DC+mirin and DC+KU-60019-treated rats (P<0.0001, one-way ANOVA (main effect)). (**D**) Mean tape sensing and removal time are restored to normal 3 weeks after treatment with mirin and KU-60019 (P<0.0001, independent sample t-test, no *post-hoc* test (DC+vehicle *vs* DC+mirin/DC+KU-60019 at 3 weeks) whilst a significant deficit remains in DC+vehicle-treated rats (# = P<0.00012, generalized linear mixed models over the whole 6 weeks). (**E**) Mean error ratio to show the number of slips *vs.* total number of steps in the horizontal ladder walking test also returns to normal 3 weeks after treatment with mirin and KU-60019 (P<0.0001, independent sample t-test, no *post-hoc* test (DC+vehicle *vs* DC+mirin/DC+KU-60019 at 3 weeks)), with a deficit remaining in DC+vehicle-treated rats (## = P<0.00011, linear mixed models over the whole 6 weeks). n = 6 rats/treatment/test, 3 independent repeats (total n = 18 rats/treatment/test).

Finally, we confirmed the results obtained with the inhibitors using specifically designed shRNA plasmids targeting Mre11 (shMre11) and ATM (shATM). These shRNA plasmids were delivered using a non-viral gene delivery vector, *in vivo*-JetPEI (PEI), which we have shown to be as efficient as AAV8 in transducing DRGN after Intra-DRG injection *in vivo and* does not invoke an off-target immune response (Almutiri *et al.*, 2018). We initially demonstrated significant knockdown of Mre11 and ATM mRNA (84% compared to DC+PEI-shControl) *in vivo* using *in vivo*-JetPEI (Supplementary Fig. 4A). The intensity and the number of γH2Ax^+^ pixels/cell as a measure of double strand breaks, were also significantly attenuated after DC+PEI-shMre11 and DC+PEI-shATM treatment indicating that knockdown of Mre11 and ATM reduces double strand breaks (Supplementary Fig. 4B and C). Electrophysiology across the lesion site after treatment with shMre11 and shATM demonstrated similar improvements in CAP amplitude (Supplementary Fig. 4D) and CAP area (Supplementary Fig. 4E) as observed after treatment with mirin and KU-60019. The mean sensing and removal time (Supplementary Fig. 4F) and mean ratio of slips to total steps (Supplementary Fig. 4G) in the sensory and locomotor tests were also similarly improved, with animals also being indistinguishable from sham-treated animals by 3 weeks, when compared to mirin and KU-60019 treatment. These results demonstrate that knockdown of Mre11 and ATM as well as mirin and KU-60019 treatment to inhibit the DNA damage response is functionally beneficial after spinal cord injury.

## Discussion

Double strand breaks occur in both long-term and acute forms of neurological disease, including Alzheimer’s, Parkinson’s and Huntington’s diseases, amyotrophic lateral sclerosis, post-cerebral ischaemia and following spinal cord injury: potentially they are a universal feature. If double strand breaks accumulate in neurological conditions, attenuating the DNA damage response as a therapeutic strategy logically should enhance disease rather than suppress it. Here, we demonstrate the opposite is true: inhibiting apical components of the DNA damage response to mute the response to double strand breaks is beneficial in models of neurodegeneration and results in unprecedented survival, axon regeneration and recovery of function after neurotrauma.

### Attenuating the DNA damage response in neurodegenerative disease

The incidence of double strand breaks in the brains of AD patients has only recently been quantified. Double strand breaks accumulate in early-stage AD and correlate with reduced cognitive scores (Simpson *et al.*, 2015). It is plausible that these un-repaired double strand breaks trigger the cell-cycle re-entry phenomena in AD brains described in the 1990s (McShea *et al.*, 1997; Nagy *et al.*, 1997) and the increase in senescence identified more recently (He *et al.*, 2013; Wei *et al.*, 2016). Consistent with cell cycle re-entry affecting neural health and acting as trigger for apoptosis, inhibitors of cell cycle progression have shown some efficacy in cerebral ischemia models (Osuga *et al.*, 2000) and have been suggested as therapies in neurodegeneration (reviewed in (Kruman and Schwartz, 2008)). Similarly caffeine, a non-specific inhibitor of ATM, is neuroprotective against etoposide-induced DNA damage to neurons *in vitro* (Kruman *et al.*, 2004) and reducing ATM gene dosage is neuroprotective in a mouse model of HD (Lu *et al.*, 2014). However, ATM inhibitors have not yet been tested in trials for neurodegeneration. In addition to double strand breaks, ATM is activated by multiple inputs, including reactive oxygen species, has a reported 700 potential downstream phosphorylation targets in multiple pathways. In contrast, the MRN complex has no reported functions other than in DNA damage repair, which potentially makes it a simpler target for therapy.

Would long-term targeting of the MRN complex as a therapeutic strategy for neurodegeneration cause cancer? Potentially, yes. However, heterozygous carriers of mutations in *Mre11*, *Rad50*, *Nbs1*/*Nbn* have only a small increase in cancer incidence over the lifecourse (Bartkova *et al.*, 2008; Kuusisto *et al.*, 2011; Damiola *et al.*, 2014). This suggests that targeting the MRN complex may a feasible strategy, particular for more rapidly progressing neurodegenerative disorders such as ALS. Potentially, a genetic approach to partially knockdown MRN complex levels *via* shRNA would be more effective and could be largely restricted to the CNS using viral delivery.

### Attenuating the DNA damage response in optic nerve injury

Clinically, the optic nerve crush injury model is directly relevant to glaucoma and other eye diseases where RGC death occurs such as ischemic optic neuropathy. For example, we previously evaluated the effect of an siRNA to caspase-2 (siCASP2) in protecting RGC from death in the same model used in this paper (Ahmed *et al.*, 2011; Vigneswara *et al.*, 2014). The siCASP2 is currently in Phase III clinical trials for the treatment of non-arteritic ischemic optic neuropathy as a result of our preclinical work (Ahmed *et al.*, 2011); http://www.eyeactnow.com). Attenuation of the DNA damage response after optic nerve crush injury *in vivo* using Mre11 or ATM inhibitors significantly neuroprotected RGC from death. RGC neuroprotection was >90% after inhibition of mirin and KU-60019, which is on a par with the inhibition of caspase-2 using an siRNA (siCASP2) (Ahmed *et al.*, 2011; Vigneswara and Ahmed, 2016). Out of all of the strategies to promote RGC neuroprotection, siCASP2 treatment appears to be the most effective (Thomas *et al.*, 2017). Other treatments such as lens injury and cAMP+oncomudulin only protect to a maximum of 24% of RGC at 21 days after ONC (Leon *et al.*, 2000; Yin *et al.*, 2006). Even combinatorial treatments such as PTEN deletion + zymosan + cAMP only protected a maximum of 36% of RGC at 10-12 weeks after ONC (de Lima *et al.*, 2012). The delivery of CNTF, a well-studied neurotrophic factor for RGC, promoted <60% RGC survival at 2 weeks (Pernet *et al.*, 2013), declining to 31% at 5 weeks (Leaver *et al.*, 2006) and 20% at 8 weeks (Pernet *et al.*, 2013) after ONC. In contrast, twice weekly injections of mirin and KU-60019 neuroprotected >90% of RGC at 24 days after ONC. Although we have not studied time-points beyond 24 days, we would expect that this level of neuroprotection would continue as we have observed for siCASP2 for 12 weeks after ONC (Vigneswara and Ahmed, 2016). Therefore, inhibition of Mre11 and ATM to our knowledge is on par with the current best treatment for RGC neuroprotection, namely siCASP2.

Importantly, although siCASP2 offers >90% RGC neuroprotection, it does not promote RGC axon regeneration (Vigneswara *et al.*, 2014). In contrast, Mre11 and ATM inhibitors not only promoted significant RGC survival, but also promoted significant RGC axon regeneration. Other treatments such lens injury and cAMP+oncomodulin only promote the regeneration of <1000 RGC axons at 1mm from the lesion at 21 days after ONC (Leon *et al.*, 2000; Yin *et al.*, 2006). Zymosan+cAMP and PTEN deletion+Zymosan+cAMP only promote the regeneration of <100 axons at 3mm from the lesion site at 2 weeks after ONC (Kurimoto *et al.*, 2010; de Lima *et al.*, 2012). Meanwhile, CNTF only promotes the regeneration of <20 axons at 1.5mm beyond the lesion site after 2 weeks and <50 axons at 4mm beyond the lesion site after 8 weeks (Leaver *et al.*, 2006; Pernet *et al.*, 2013). Our results showing robust RGC axon regeneration of >1000 axons at 4mm beyond the lesion site suggest that inhibition of Mre11 and ATM supersedes some of the best treatments that have been reported to date to promote RGC axon regeneration. Moreover, our results demonstrate that mirin or KU-60019 on their own cannot only protect RGC from death but also promote their axons to regenerate, making translation to the clinical scenario easier due to the requirement of delivering a single molecule to activate both neuroprotective and axon regenerative pathways.

### Attenuating the DNA damage response in SCI

Attenuation of the DNA damage response after DC injury using Mre11 or ATM inhibitors promoted significant DC axon regeneration and electrophysiological, sensory and locomotor improvements. This is despite the presence of spinal cord cavities that are characteristic of DC crush injury in the rat. However, after DC injury, DRGN do not die and hence neuroprotection *in vivo* could not be assessed. Nonetheless, *in vitro* experiments with both Mre11 and ATM inhibitors demonstrated significant DRGN neuroprotection and hence we can surmise that the inhibitors are DRGN neuroprotective. The recovery of function after SCI in adult rats administered Mre11 or ATM inhibitors and confirmed using shMre11 and shATM, is striking and, to the best of our knowledge, unprecedented. The improvements in sensory and locomotor function that we achieved are perhaps more remarkable given that DC injury in adult rats results in spinal cord cavities but cavitation is absent after DC injury in the mouse (Surey *et al.*, 2014). Instead in the mouse, the lesion sites fill with dense fibrous connective tissues which allow some regeneration of axons (Bilgen *et al.*, 2007). In contrast, cavitation in the rat results in worsening of the initial injury, leading to disconnection of more axons over a period of several months (Tang *et al.*, 2003; Byrnes *et al.*, 2010).

Blunt injury, such as compression or contusion, account for more than >80% of all SCI in humans (Taoka and Okajima, 1998; Metz *et al.*, 2000; Brechtel *et al.*, 2006). Our rat DC crush model replicates many aspects of the human pathology well, including evolution of the injury over time and morphological aspects, such as spinal cord atrophy, myelomalacia, cavity, cyst and syrinx formation and cord disruption (Gruner, 1992; Taoka and Okajima, 1998; Kwon *et al.*, 2002). Importantly, cavity and cysts form and increase over time, thus transecting more axons and causing ongoing damage to the spinal cord architecture. Our study, therefore, addresses axon regeneration and functional recovery in a model that cavitates, as opposed to mouse models where cavitation is absent. Taken together, these clinically relevant features suggest our approach could be a very effective therapy for patients.

The Mre11 and ATM inhibitors used in this study are potent, small molecule inhibitors that are specific for their targets and have a low IC_50_. These inhibitors were developed has chemotherapy agents for cancer. DNA damage-induced tumour cell killing has been demonstrated in preclinical models with mirin and KU-60019 (Hosoya and Miyagawa, 2014) and other ATM inhibitors are currently in trials that potentially could be re-purposed for SCI. Importantly, our method of administering the compounds – by intrathecal injection – can be translated directly to human therapy, This method gives direct access of compounds to the injury site, which often requires lower drug doses with greater efficiency than other routes of administration (Bottros and Christo, 2014), limiting the potential unwanted side effects on other cells in the CNS such as glia.

In our experiments, the mirin and KU-60019 compounds were delivered directly after injury but this could also happen in human patients since most cases attend emergency care immediately. For neurodegeneration, a viral-mediated genetic knockdown approach may prove to be more effective by effecting a partial knockdown in MRN complex levels over a longer period; this may also prove to be the case in SCI treatments. However, there are caveats to this approach. Inhibiting the DNA damage response may have negative consequences to cells and system functions if applied systemically. Even if used intrathecally as suggested by us, there may be negative consequences on dividing cells such as glia in the CNS. Hence, caution must be used when using these compounds in treating human conditions.

Current therapies for SCI are limited and only offer palliative relief with none reversing the underlying damage to axons. Methylprednisolone is the only pharmacotherapy that is currently approved for use in SCI, but this has failed to show clinically significant effects and is now often used off-label (Varma *et al.*, 2013). Several other experimental therapies are currently in the early phases of development, including the Rho antagonist, Cethrin; anti-Nogo antibodies and autologous mesenchymal stem cell transplants. However, completed clinical trials of these reagents have either not been reported, abandoned or concluded with no significant improvements reported. Consequently, there is a clear unmet need for a therapy in SCI that would not only promote neuroprotection but also promote axon regeneration. It is clear from our previous work that neuroprotection and axon regeneration are signalled by different signalling pathways (Ahmed *et al.*, 2010) and our study is the first to report single inhibitors capable of stimulating both processes. A ‘single hit’ molecule that can modify both neuroprotective and axon regenerative pathways would be an additional advantage for a potential therapy to treat the devastating loss of function that ensues after SCI in humans. Here, we report that DNA damage response inhibitors are good candidates.

In conclusion, our paper highlights a novel therapeutic strategy by targeting the response of neurons to double strand breaks to protect neurons from death and promote their axon regeneration. This strategy prevents loss of function in multiple neuropathological paradigms and suggests new treatment possibilities for neurological conditions.

## Acknowledgments

We thank Emma Richards for help with the initial stages of this project, Drs Damien Crowther and Mel Feany for providing *Drosophila* stocks and Professors Martin Berry and Malcolm Taylor for helpful comments on the manuscript.

## Supplementary Figures

**Supplementary Figure 1.** Image processing of DNA double strand breaks in *Drosophila* neurons. 41 × 41 × 13.5 μm volumes of adult central brain neurons stained with anti-pH2Av were visualized by confocal microscopy with 0.45 m step sizes in z. A 63x N.A. 1.2 water immersion lens was used mounted on an inverted Zeiss LSM880 microscope and the Airy scan module used with super-resolution settings to improve resolution. Anti-pH2Av shows a uniform nuclear fluorescence both in our hands and in a previous report (Bellesi *et al.*, 2016). Left panels show the projected z-series of images without adjustments to contrast or brightness. The right panels are the identical images processed to remove the low intensity fluorescence by resetting the black level to display only the bright pH2Av^+^ foci. Control neurons have a single pH2Av^+^ focus corresponding to the nucleolus organizing region. Arrowheads point to clusters of foci in neurons expressing Aβ_1-42_. Scale bars = 10µm.

**Supplementary Figure 2.** Genetic depletion of the MRN complex is neuroprotective in *Drosophila* neurodegeneration models. (**A**) Quantification of Tau.R406W expression in the heads of flies by Western blot. Flies were developed at 18 °C to inhibit expression of Tau then decapitated 0, 7, 14 or 21 d after shifting to 29 °C. Tau proteins levels are not altered in *nbs^−/+^* heterozygotes. Phosphorylation of Tau at Thr181 is also unchanged. Quantification of Tau levels normalized to actin and the Tau:pTau ratio is shown below (n = 3; mean ± SEM; ANOVA with Dunn’s post-hoc test). (**B**) Negative geotaxis climbing assay. Flies expressing Htt.Q128 show premature decline in climbing ability. This is partially suppressed in flies heterozygous for a null *rad50^EP1^* allele (p=0.0372) despite *rad50* heterozygous flies already showing reduced climbing ability *vs*. the *w^1118^* control (p=0.0007). Lines were fitted using non-linear regression and compared by extra sum-of-squares F-test. (**C**) Expression of Htt.Q128 in the eye leads to loss of pigment-containing cells in the retina after 7 weeks. Pigment loss is largely prevented in flies *rad50^−/+^* flies. (**D**) and (**E**) Negative geotaxis climbing assays of flies expressing two forms of tandem Aβ_1-42_ expression with no flexible linker or with a 12-amino acid flexible linker (**D**) in adult neurons (Speretta *et al.*, 2012). Flies climbed in vials with three marked zones. Aβ_1-42_ expressing (red) flies were compared to Aβ_1-42_; *nbs^1/+^* flies (blue) on the days indicated after shifting to 29 °C to induce expression. Comparison was by χ^2^ test (* indicates P<0.05). (**F**) Circadian periodicity analyzed by cosinor analysis shows similar results to the CLEAN spectral analysis shown in Fig. 1F (Rosato and Kyriacou, 2006). Comparison by ANOVA with Tukey’s post-hoc test; p values and n are indicated.

**Methods for Tau quantification in (A).** 15 adult flies per genotype and time point were decapitated with a razor blade then homogenized in 150 μl RIPA buffer (New England Biolabs) supplemented with protease and phosphatase inhibitor cocktails (both Merck). Homogenization used a MP Biomedicals FastPrep blender running at 6.0 m/s for 20 s in screwcap tubes containing matrix C garnet beads (MP Biomedicals). Lysates were clarified by centrifugation at 12000 × g for 5 min at 4 °C then 6X Laemlli buffer + 50 mM DTT was added and the samples boiled for 5 min. Proteins were separated on 12% Tris.Glycine SDS-PAGE gels and transferred to PVDF membranes. Primary antibodies used were: rabbit anti-total Tau (1:1.000; DAKO); mouse anti-Tau pThr181 (1:500; ThermoFisher) and mouse IgM anti-actin JLA20 (1:5.000; Developmental Studies Hybridoma Bank). Secondary antibodies used were: HRP-conjugated horse anti-mouse IgG and HRP-conjugated horse anti-rabbit IgG (both 1:10,000; New England Biolabs; note: the anti-mouse IgG secondary antibody cross-reacts with mouse IgM). Membranes were visualized using a Vilber Fusion FX scanner. Total Tau levels were normalized to actin for each sample and phosphorylated Tau normalized to total Tau using ImageJ.

**Supplementary Figure 3.** Inhibition of Mre11 prevents DRGN apoptosis and stimulates neurite outgrowth in the presence of inhibitory CME. (**A**) Adult DRGN stained at 4 days after plating, with anti-γH2Ax antibodies to demonstrate double strand breaks (arrows) in DRGN *in vitro*, whilst satellite glial cells did not possess double strand breaks (arrowheads). (**B**) Representative images from DRGN after 4 days in culture and treated with vehicle, FGF2 (positive control), mirin and KU-60019. (**C-E**) Mirin or KU-60019 treatment enhances (**C**), DRGN survival, (**D**) % DRGN with neurites and (**E**) increased the mean length of the longest neurite. Comparison in (C-E) by one-way ANOVA with Dunnett’s post-hoc test, n = 3 wells/condition, 3 independent repeats (total n = 9 wells/condition (all 9 data points shown on graphs)). Scale bars in **A** = 100µm and **B** = 50µm.

**Supplementary Figure 4.** Intra-DRG injection of DNA plasmids to knock down Mre11 and ATM using non-viral *in vivo*-JetPEI (PEI) promotes significant functional repair after DC injury *in vivo*. (**A**) PEI delivered plasmids significantly knock down Mre11 and ATM mRNA expression in spinal L4/L5 DRGs at 4 weeks after DC injury. (**B**) Immunohistochemistry for γH2Ax to demonstrate DNA breaks 4 weeks after DC injury and their attenuation by shMre11 and shATM in DRGN. (**C**) Quantification of γH2Ax+ pixels/cells to demonstrate significant attenuation of double strand breaks by shMre11 and shATM. (**D**) Negative CAP amplitudes were significantly attenuated in DC+vehicle-treated rats but were significantly improved in DC+PEI-shMre11 and DC+PEI-shATM treatment (P<0.0001, one-way ANOVA (main effect)). (**E**) Mean CAP area at different stimulation intensities were significantly attenuated in DC+vehicle-treated rats but improved significantly in DC+PEI-shMre11 and DC+PEI-shATM-treated rats (P<0.0001, one-way ANOVA (main effect)). (**F**) Mean tape sensing and removal times are restored to normal 3 weeks after treatment with PEI-shMre11 and PEI-shATM (P<0.0001, independent sample t-test (DC+vehicle vs DC+PEI-shMre11/DC+PEI-shATM at 3 weeks) whilst a significant deficit remains in DC+vehicle-treated rats (# = P<0.00011, generalized linear mixed models over the whole 6 weeks). (**G**) Mean error ratio to show the number of slips vs. total number of steps in the horizontal ladder walking test also returns to normal 3 weeks after treatment with PEI-shMre11 and PEI-shATM (P<0.0001, independent sample t-test (DC+vehicle vs DC+PEI-shMre11/DC+PEI-shATM at 3 weeks)), with a deficit remaining in DC+vehicle-treated rats (## = P<0.00014, linear mixed models over the whole 6 weeks). AU = arbitrary units. n = 6 rats/treatment/test, 3 independent repeats (total n = 18 rats/treatment/test). Scale bars in **B** = 50µm, insets in **B** = 10µm.

## Movies

**Movie 1.** Rotated views of 3D rendered volumes showing pH2Av^+^ foci in the nuclei of central brain neurons from control and Aβ_1-42_ expressing adult flies. Volumes shown in Fig. 1 and Supplementary Fig. 1 were rendered in Fiji-3. Control brain is to the left and Aβ_1-42_-expressing brain to the right.

**Movie 2.** Genetic reduction of *nbs* levels maintains neural activity in flies after 10 d of A expression. Flies expressing tandem Aβ_1-42_ (Speretta *et al.*, 2012) specifically in adult neurons for 10 d under the control of Elav-Gal4. *Gal80^ts^* was included to prevent expression until eclosion. Flies were maintained at 18 °C during development and shifted to the permissive temperature of 29°C upon eclosion. Flies in the vial to the left are wild-type for *nbs;* flies to the right are heterozygous for the null *nbs^2^* allele.

## References

Ahmed Z, Aslam M, Lorber B, Suggate EL, Berry M, Logan A. Optic nerve and vitreal inflammation are both RGC neuroprotective but only the latter is RGC axogenic. Neurobiol Dis 2010; 37(2): 441–54.

Ahmed Z, Bansal D, Tizzard K, Surey S, Esmaeili M, Gonzalez AM, et al. Decorin blocks scarring and cystic cavitation in acute and induces scar dissolution in chronic spinal cord wounds. Neurobiol Dis 2014; 64: 163–76.

Ahmed Z, Dent RG, Suggate EL, Barrett LB, Seabright RJ, Berry M, et al. Disinhibition of neurotrophin-induced dorsal root ganglion cell neurite outgrowth on CNS myelin by siRNA-mediated knockdown of NgR, p75NTR and Rho-A. Mol Cell Neurosci 2005; 28(3): 509–23.

Ahmed Z, Kalinski H, Berry M, Almasieh M, Ashush H, Slager N, et al. Ocular neuroprotection by siRNA targeting caspase-2. Cell Death Dis 2011; 2: e173.

Ahmed Z, Mazibrada G, Seabright RJ, Dent RG, Berry M, Logan A. TACE-induced cleavage of NgR and p75NTR in dorsal root ganglion cultures disinhibits outgrowth and promotes branching of neurites in the presence of inhibitory CNS myelin. FASEB J 2006a; 20(11): 1939–41.

Ahmed Z, Suggate EL, Brown ER, Dent RG, Armstrong SJ, Barrett LB, et al. Schwann cell-derived factor-induced modulation of the NgR/p75NTR/EGFR axis disinhibits axon growth through CNS myelin in vivo and in vitro. Brain 2006b; 129(Pt 6): 1517–33.

Almutiri S, Berry M, Logan A, Ahmed Z. Non-viral-mediated suppression of AMIGO3 promotes disinhibited NT3-mediated regeneration of spinal cord dorsal column axons. Sci Rep 2018; 8(1): 10707.

Bartkova J, Tommiska J, Oplustilova L, Aaltonen K, Tamminen A, Heikkinen T, et al. Aberrations of the MRE11-RAD50-NBS1 DNA damage sensor complex in human breast cancer: MRE11 as a candidate familial cancer-predisposing gene. Mol Oncol 2008; 2(4): 296–316.

Bellesi M, Bushey D, Chini M, Tononi G, Cirelli C. Contribution of sleep to the repair of neuronal DNA double-strand breaks: evidence from flies and mice. Sci Rep 2016; 6: 36804.

Berry M, Carlile J, Hunter A. Peripheral nerve explants grafted into the vitreous body of the eye promote the regeneration of retinal ganglion cell axons severed in the optic nerve. J Neurocytol 1996; 25(2): 147–70.

Bilgen M, Al-Hafez B, Alrefae T, He YY, Smirnova IV, Aldur MM, et al. Longitudinal magnetic resonance imaging of spinal cord injury in mouse: changes in signal patterns associated with the inflammatory response. Magn Reson Imaging 2007; 25(5): 657–64.

Bottros MM, Christo PJ. Current perspectives on intrathecal drug delivery. J Pain Res 2014; 7: 615–26.

Brechtel K, Tura A, Abdibzadeh M, Hirsch S, Conrad S, Schwab JM. Intrinsic locomotor outcome in dorsal transection of rat spinal cord: predictive value of minimal incision depth. Spinal Cord 2006; 44(10): 605–13.

Byrnes KR, Fricke ST, Faden AI. Neuropathological differences between rats and mice after spinal cord injury. J Magn Reson Imaging 2010; 32(4): 836–46.

Celeste A, Petersen S, Romanienko PJ, Fernandez-Capetillo O, Chen HT, Sedelnikova OA, et al. Genomic instability in mice lacking histone H2AX. Science 2002; 296(5569): 922–7.

Damiola F, Pertesi M, Oliver J, Le Calvez-Kelm F, Voegele C, Young EL, et al. Rare key functional domain missense substitutions in MRE11A, RAD50, and NBN contribute to breast cancer susceptibility: results from a Breast Cancer Family Registry case-control mutation-screening study. Breast Cancer Res 2014; 16(3): R58.

de Lima S, Koriyama Y, Kurimoto T, Oliveira JT, Yin Y, Li Y, et al. Full-length axon regeneration in the adult mouse optic nerve and partial recovery of simple visual behaviors. Proc Natl Acad Sci U S A 2012; 109(23): 9149–54.

Dissel S, Codd V, Fedic R, Garner KJ, Costa R, Kyriacou CP, et al. A constitutively active cryptochrome in Drosophila melanogaster. Nat Neurosci 2004; 7(8): 834–40.

Douglas MR, Morrison KC, Jacques SJ, Leadbeater WE, Gonzalez AM, Berry M, et al. Off-target effects of epidermal growth factor receptor antagonists mediate retinal ganglion cell disinhibited axon growth. Brain 2009; 132(Pt 11): 3102–21.

Dupre A, Boyer-Chatenet L, Sattler RM, Modi AP, Lee JH, Nicolette ML, et al. A forward chemical genetic screen reveals an inhibitor of the Mre11-Rad50-Nbs1 complex. Nat Chem Biol 2008; 4(2): 119–25.

Fagoe ND, Attwell CL, Eggers R, Tuinenbreijer L, Kouwenhoven D, Verhaagen J, et al. Evaluation of Five Tests for Sensitivity to Functional Deficits following Cervical or Thoracic Dorsal Column Transection in the Rat. PLoS One 2016; 11(3): e0150141.

Farrukh F, Davies E, Berry M, Logan A, Ahmed Z. BMP4/Smad1 Signalling Promotes Spinal Dorsal Column Axon Regeneration and Functional Recovery After Injury. Mol Neurobiol 2019.

Faville R, Kottler B, Goodhill GJ, Shaw PJ, van Swinderen B. How deeply does your mutant sleep? Probing arousal to better understand sleep defects in Drosophila. Sci Rep 2015; 5: 8454.

Fielder E, von Zglinicki T, Jurk D. The DNA Damage Response in Neurons: Die by Apoptosis or Survive in a Senescence-Like State? J Alzheimers Dis 2017; 60(s1): S107–S31.

Garner KM, Pletnev AA, Eastman A. Corrected structure of mirin, a small-molecule inhibitor of the Mre11-Rad50-Nbs1 complex. Nat Chem Biol 2009; 5(3): 129–30; author reply 30.

Gillingwater TH, Wishart TM. Mechanisms underlying synaptic vulnerability and degeneration in neurodegenerative disease. Neuropathol Appl Neurobiol 2013; 39(4): 320–34.

Golding SE, Rosenberg E, Valerie N, Hussaini I, Frigerio M, Cockcroft XF, et al. Improved ATM kinase inhibitor KU-60019 radiosensitizes glioma cells, compromises insulin, AKT and ERK prosurvival signaling, and inhibits migration and invasion. Mol Cancer Ther 2009; 8(10): 2894–902.

Gruner JA. A monitored contusion model of spinal cord injury in the rat. J Neurotrauma 1992; 9(2): 123–6; discussion 6-8.

Hains BC, Saab CY, Lo AC, Waxman SG. Sodium channel blockade with phenytoin protects spinal cord axons, enhances axonal conduction, and improves functional motor recovery after contusion SCI. Exp Neurol 2004; 188(2): 365–77.

Hata K, Fujitani M, Yasuda Y, Doya H, Saito T, Yamagishi S, et al. RGMa inhibition promotes axonal growth and recovery after spinal cord injury. J Cell Biol 2006; 173(1): 47–58.

Hayashi T, Sakurai M, Abe K, Sadahiro M, Tabayashi K, Itoyama Y. Apoptosis of motor neurons with induction of caspases in the spinal cord after ischemia. Stroke 1998; 29(5): 1007–12; discussion 13.

He M, Di GH, Cao AY, Hu Z, Jin W, Shen ZZ, et al. RAD50 and NBS1 are not likely to be susceptibility genes in Chinese non-BRCA1/2 hereditary breast cancer. Breast Cancer Res Treat 2012; 133(1): 111–6.

He N, Jin WL, Lok KH, Wang Y, Yin M, Wang ZJ. Amyloid-beta(1-42) oligomer accelerates senescence in adult hippocampal neural stem/progenitor cells via formylpeptide receptor 2. Cell Death Dis 2013; 4: e924.

Herrup K, Neve R, Ackerman SL, Copani A. Divide and die: cell cycle events as triggers of nerve cell death. J Neurosci 2004; 24(42): 9232–9.

Hosoya N, Miyagawa K. Targeting DNA damage response in cancer therapy. Cancer Sci 2014; 105(4): 370–88.

Jacques SJ, Ahmed Z, Forbes A, Douglas MR, Vigenswara V, Berry M, et al. AAV8(gfp) preferentially targets large diameter dorsal root ganglion neurones after both intra-dorsal root ganglion and intrathecal injection. Mol Cell Neurosci 2012; 49(4): 464–74.

Katchanov J, Harms C, Gertz K, Hauck L, Waeber C, Hirt L, et al. Mild cerebral ischemia induces loss of cyclin-dependent kinase inhibitors and activation of cell cycle machinery before delayed neuronal cell death. J Neurosci 2001; 21(14): 5045–53.

Kerr F, Augustin H, Piper MD, Gandy C, Allen MJ, Lovestone S, et al. Dietary restriction delays aging, but not neuronal dysfunction, in Drosophila models of Alzheimer’s disease. Neurobiol Aging 2011; 32(11): 1977–89.

Kotipatruni RR, Dasari VR, Veeravalli KK, Dinh DH, Fassett D, Rao JS. p53- and Bax-mediated apoptosis in injured rat spinal cord. Neurochem Res 2011; 36(11): 2063–74.

Kruman, II, Schwartz EI. DNA damage response and neuroprotection. Front Biosci 2008; 13: 2504–15.

Kruman, II, Wersto RP, Cardozo-Pelaez F, Smilenov L, Chan SL, Chrest FJ, et al. Cell cycle activation linked to neuronal cell death initiated by DNA damage. Neuron 2004; 41(4): 549–61.

Kurimoto T, Yin Y, Omura K, Gilbert HY, Kim D, Cen LP, et al. Long-distance axon regeneration in the mature optic nerve: contributions of oncomodulin, cAMP, and pten gene deletion. J Neurosci 2010; 30(46): 15654–63.

Kuusisto KM, Bebel A, Vihinen M, Schleutker J, Sallinen SL. Screening for BRCA1, BRCA2, CHEK2, PALB2, BRIP1, RAD50, and CDH1 mutations in high-risk Finnish BRCA1/2-founder mutation-negative breast and/or ovarian cancer individuals. Breast Cancer Res 2011; 13(1): R20.

Kwon BK, Oxland TR, Tetzlaff W. Animal models used in spinal cord regeneration research. Spine (Phila Pa 1976) 2002; 27(14): 1504–10.

Lamarche BJ, Orazio NI, Weitzman MD. The MRN complex in double-strand break repair and telomere maintenance. FEBS Lett 2010; 584(17): 3682–95.

Leaver SG, Cui Q, Bernard O, Harvey AR. Cooperative effects of bcl-2 and AAV-mediated expression of CNTF on retinal ganglion cell survival and axonal regeneration in adult transgenic mice. Eur J Neurosci 2006; 24(12): 3323–32.

Lee JH, Paull TT. ATM activation by DNA double-strand breaks through the Mre11-Rad50-Nbs1 complex. Science 2005; 308(5721): 551–4.

Leon S, Yin Y, Nguyen J, Irwin N, Benowitz LI. Lens injury stimulates axon regeneration in the mature rat optic nerve. J Neurosci 2000; 20(12): 4615–26.

Levine JD, Funes P, Dowse HB, Hall JC. Advanced analysis of a cryptochrome mutation’s effects on the robustness and phase of molecular cycles in isolated peripheral tissues of Drosophila. BMC Neurosci 2002; 3: 5.

Lo AC, Saab CY, Black JA, Waxman SG. Phenytoin protects spinal cord axons and preserves axonal conduction and neurological function in a model of neuroinflammation in vivo. J Neurophysiol 2003; 90(5): 3566–71.

Long DM, Blake MR, Dutta S, Holbrook SD, Kotwica-Rolinska J, Kretzschmar D, et al. Relationships between the circadian system and Alzheimer’s disease-like symptoms in Drosophila. PLoS One 2014; 9(8): e106068.

Love S. Neuronal expression of cell cycle-related proteins after brain ischaemia in man. Neurosci Lett 2003; 353(1): 29–32.

Lu XH, Mattis VB, Wang N, Al-Ramahi I, van den Berg N, Fratantoni SA, et al. Targeting ATM ameliorates mutant Huntingtin toxicity in cell and animal models of Huntington’s disease. Sci Transl Med 2014; 6(268): 268ra178.

Marsh J, Bagol SH, Williams RSB, Dickson G, Alifragis P. Synapsin I phosphorylation is dysregulated by beta-amyloid oligomers and restored by valproic acid. Neurobiol Dis 2017; 106: 63–75.

McShea A, Harris PL, Webster KR, Wahl AF, Smith MA. Abnormal expression of the cell cycle regulators P16 and CDK4 in Alzheimer’s disease. Am J Pathol 1997; 150(6): 1933–9.

Merlo D, Mollinari C, Racaniello M, Garaci E, Cardinale A. DNA Double Strand Breaks: A Common Theme in Neurodegenerative Diseases. Curr Alzheimer Res 2016; 13(11): 1208–18.

Metz GA, Curt A, van de Meent H, Klusman I, Schwab ME, Dietz V. Validation of the weight-drop contusion model in rats: a comparative study of human spinal cord injury. J Neurotrauma 2000; 17(1): 1–17.

Milanese C, Cerri S, Ulusoy A, Gornati SV, Plat A, Gabriels S, et al. Activation of the DNA damage response in vivo in synucleinopathy models of Parkinson’s disease. Cell Death Dis 2018; 9(8): 818.

Mladenov E, Magin S, Soni A, Iliakis G. DNA double-strand-break repair in higher eukaryotes and its role in genomic instability and cancer: Cell cycle and proliferation-dependent regulation. Semin Cancer Biol 2016; 37-38: 51–64.

Nagy Z, Esiri MM, Cato AM, Smith AD. Cell cycle markers in the hippocampus in Alzheimer’s disease. Acta Neuropathol 1997; 94(1): 6–15.

Neumann S, Bradke F, Tessier-Lavigne M, Basbaum AI. Regeneration of sensory axons within the injured spinal cord induced by intraganglionic cAMP elevation. Neuron 2002; 34(6): 885–93.

Neumann S, Woolf CJ. Regeneration of dorsal column fibers into and beyond the lesion site following adult spinal cord injury. Neuron 1999; 23(1): 83–91.

Osuga H, Osuga S, Wang F, Fetni R, Hogan MJ, Slack RS, et al. Cyclin-dependent kinases as a therapeutic target for stroke. Proc Natl Acad Sci U S A 2000; 97(18): 10254–9.

Pernet V, Joly S, Dalkara D, Jordi N, Schwarz O, Christ F, et al. Long-distance axonal regeneration induced by CNTF gene transfer is impaired by axonal misguidance in the injured adult optic nerve. Neurobiol Dis 2013; 51: 202–13.

Renn SC, Park JH, Rosbash M, Hall JC, Taghert PH. A pdf neuropeptide gene mutation and ablation of PDF neurons each cause severe abnormalities of behavioral circadian rhythms in Drosophila. Cell 1999; 99(7): 791–802.

Romero E, Cha GH, Verstreken P, Ly CV, Hughes RE, Bellen HJ, et al. Suppression of neurodegeneration and increased neurotransmission caused by expanded full-length huntingtin accumulating in the cytoplasm. Neuron 2008; 57(1): 27–40.

Rosato E, Kyriacou CP. Analysis of locomotor activity rhythms in Drosophila. Nat Protoc 2006; 1(2): 559–68.

Russell CL, Semerdjieva S, Empson RM, Austen BM, Beesley PW, Alifragis P. Amyloid-beta acts as a regulator of neurotransmitter release disrupting the interaction between synaptophysin and VAMP2. PLoS One 2012; 7(8): e43201.

Saeed Y, Abbott SM. Circadian Disruption Associated with Alzheimer’s Disease. Curr Neurol Neurosci Rep 2017; 17(4): 29.

Shiloh Y, Ziv Y. The ATM protein kinase: regulating the cellular response to genotoxic stress, and more. Nat Rev Mol Cell Biol 2013; 14(4): 197–210.

Shrivastav M, De Haro LP, Nickoloff JA. Regulation of DNA double-strand break repair pathway choice. Cell Res 2008; 18(1): 134–47.

Simpson JE, Ince PG, Matthews FE, Shaw PJ, Heath PR, Brayne C, et al. A neuronal DNA damage response is detected at the earliest stages of Alzheimer’s neuropathology and correlates with cognitive impairment in the Medical Research Council’s Cognitive Function and Ageing Study ageing brain cohort. Neuropathol Appl Neurobiol 2015; 41(4): 483–96.

Speretta E, Jahn TR, Tartaglia GG, Favrin G, Barros TP, Imarisio S, et al. Expression in drosophila of tandem amyloid beta peptides provides insights into links between aggregation and neurotoxicity. J Biol Chem 2012; 287(24): 20748–54.

Stivers JT. Small molecule versus DNA repair nanomachine. Nat Chem Biol 2008; 4(2): 86–8.

Suberbielle E, Sanchez PE, Kravitz AV, Wang X, Ho K, Eilertson K, et al. Physiologic brain activity causes DNA double-strand breaks in neurons, with exacerbation by amyloid-beta. Nat Neurosci 2013; 16(5): 613–21.

Surey S, Berry M, Logan A, Bicknell R, Ahmed Z. Differential cavitation, angiogenesis and wound-healing responses in injured mouse and rat spinal cords. Neuroscience 2014; 275: 62–80.

Tang X, Davies JE, Davies SJ. Changes in distribution, cell associations, and protein expression levels of NG2, neurocan, phosphacan, brevican, versican V2, and tenascin-C during acute to chronic maturation of spinal cord scar tissue. J Neurosci Res 2003; 71(3): 427–44.

Taoka Y, Okajima K. Spinal cord injury in the rat. Prog Neurobiol 1998; 56(3): 341–58.

Thomas CN, Berry M, Logan A, Blanch RJ, Ahmed Z. Caspases in retinal ganglion cell death and axon regeneration. Cell Death Discov 2017; 3: 17032.

Varma AK, Das A, Wallace Gt, Barry J, Vertegel AA, Ray SK, et al. Spinal cord injury: a review of current therapy, future treatments, and basic science frontiers. Neurochem Res 2013; 38(5): 895–905.

Vigneswara V, Ahmed Z. Long-term neuroprotection of retinal ganglion cells by inhibiting caspase-2. Cell Death Discov 2016; 2: 16044.

Vigneswara V, Akpan N, Berry M, Logan A, Troy CM, Ahmed Z. Combined suppression of CASP2 and CASP6 protects retinal ganglion cells from apoptosis and promotes axon regeneration through CNTF-mediated JAK/STAT signalling. Brain 2014; 137(Pt 6): 1656–75.

Vigneswara V, Berry M, Logan A, Ahmed Z. Caspase-2 is upregulated after sciatic nerve transection and its inhibition protects dorsal root ganglion neurons from apoptosis after serum withdrawal. PLoS One 2013; 8(2): e57861.

Wei Z, Chen XC, Song Y, Pan XD, Dai XM, Zhang J, et al. Amyloid beta Protein Aggravates Neuronal Senescence and Cognitive Deficits in 5XFAD Mouse Model of Alzheimer’s Disease. Chin Med J (Engl) 2016; 129(15): 1835–44.

Wen Y, Yang S, Liu R, Brun-Zinkernagel AM, Koulen P, Simpkins JW. Transient cerebral ischemia induces aberrant neuronal cell cycle re-entry and Alzheimer’s disease-like tauopathy in female rats. J Biol Chem 2004; 279(21): 22684–92.

Wittmann CW, Wszolek MF, Shulman JM, Salvaterra PM, Lewis J, Hutton M, et al. Tauopathy in Drosophila: neurodegeneration without neurofibrillary tangles. Science 2001; 293(5530): 711–4.

Yaksh TL, Rudy TA. Chronic catheterization of the spinal subarachnoid space. Physiol Behav 1976; 17(6): 1031–6.

Yang Y, Wu N, Tian S, Li F, Hu H, Chen P, et al. Lithium promotes DNA stability and survival of ischemic retinal neurocytes by upregulating DNA ligase IV. Cell Death Dis 2016; 7(11): e2473.

Yin Y, Henzl MT, Lorber B, Nakazawa T, Thomas TT, Jiang F, et al. Oncomodulin is a macrophage-derived signal for axon regeneration in retinal ganglion cells. Nat Neurosci 2006; 9(6): 843–52.

