## Supplementary data for "Attenuating the DNA damage response to double strand breaks restores function in models of CNS neurodegeneration"

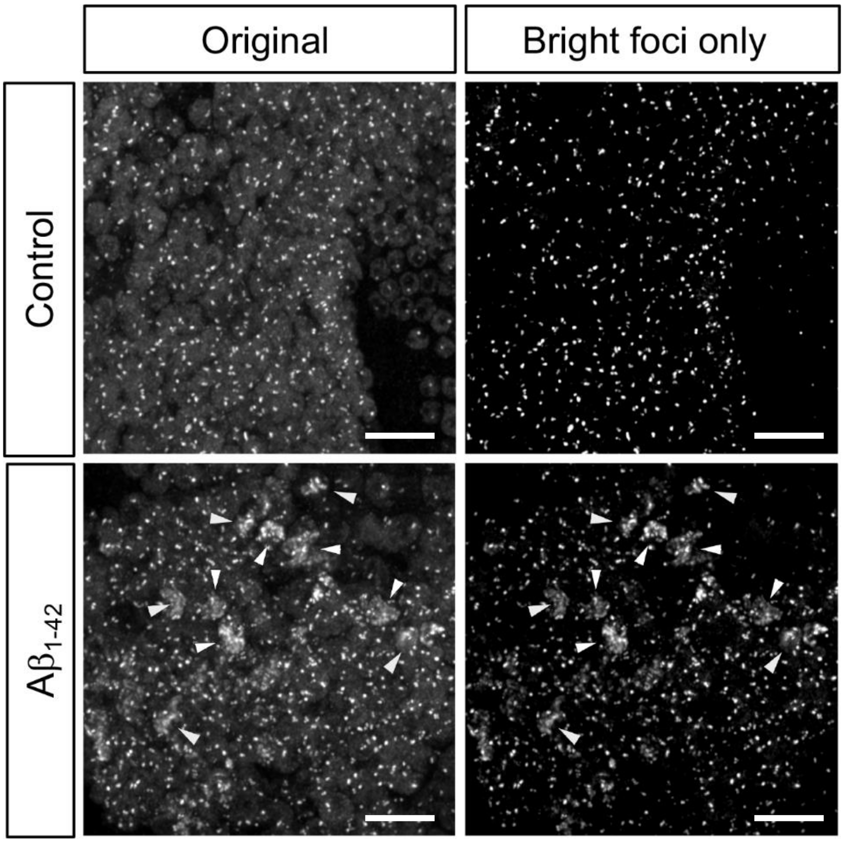


**Supplementary Fig. 1**

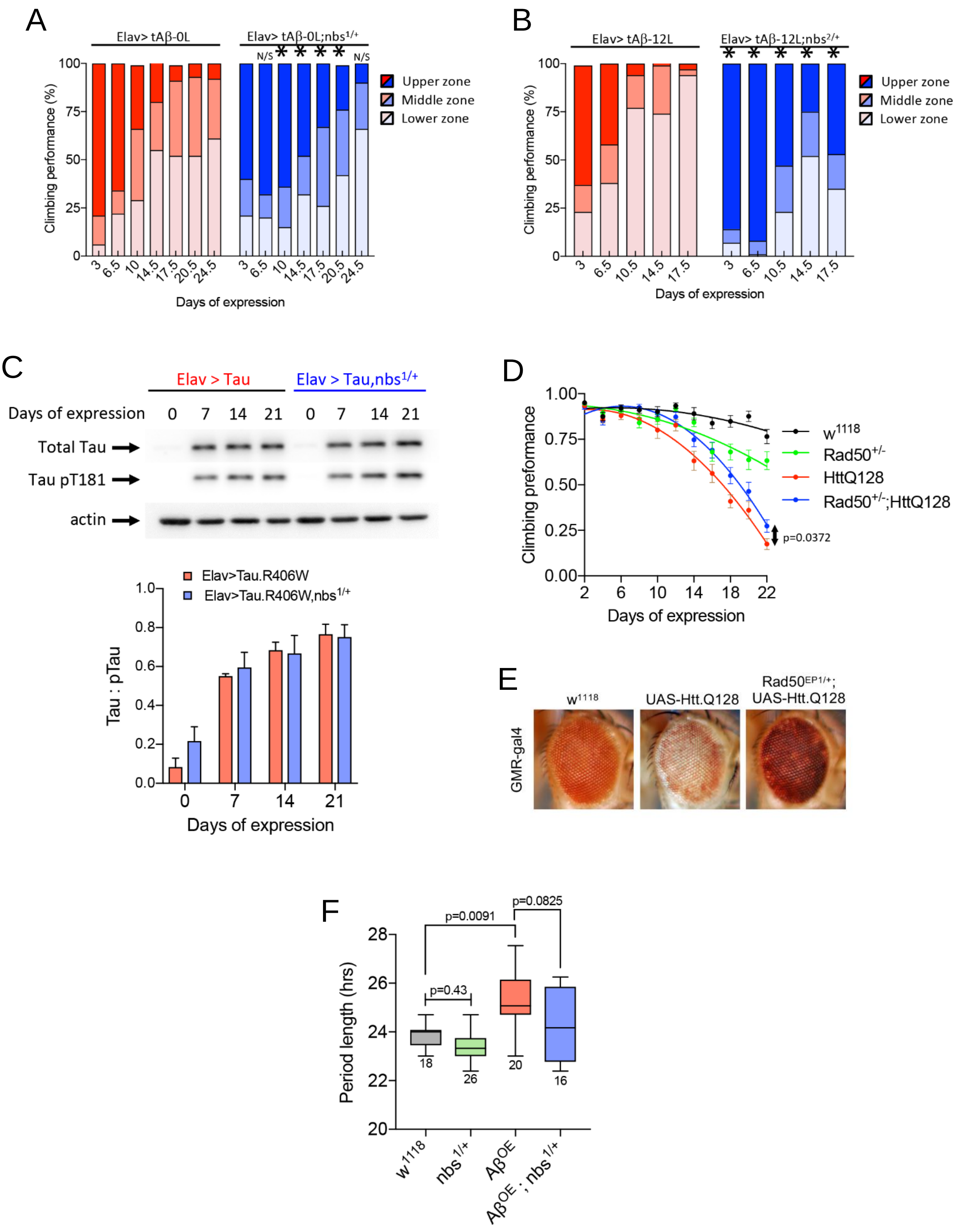


**Supplementary Fig. 2**

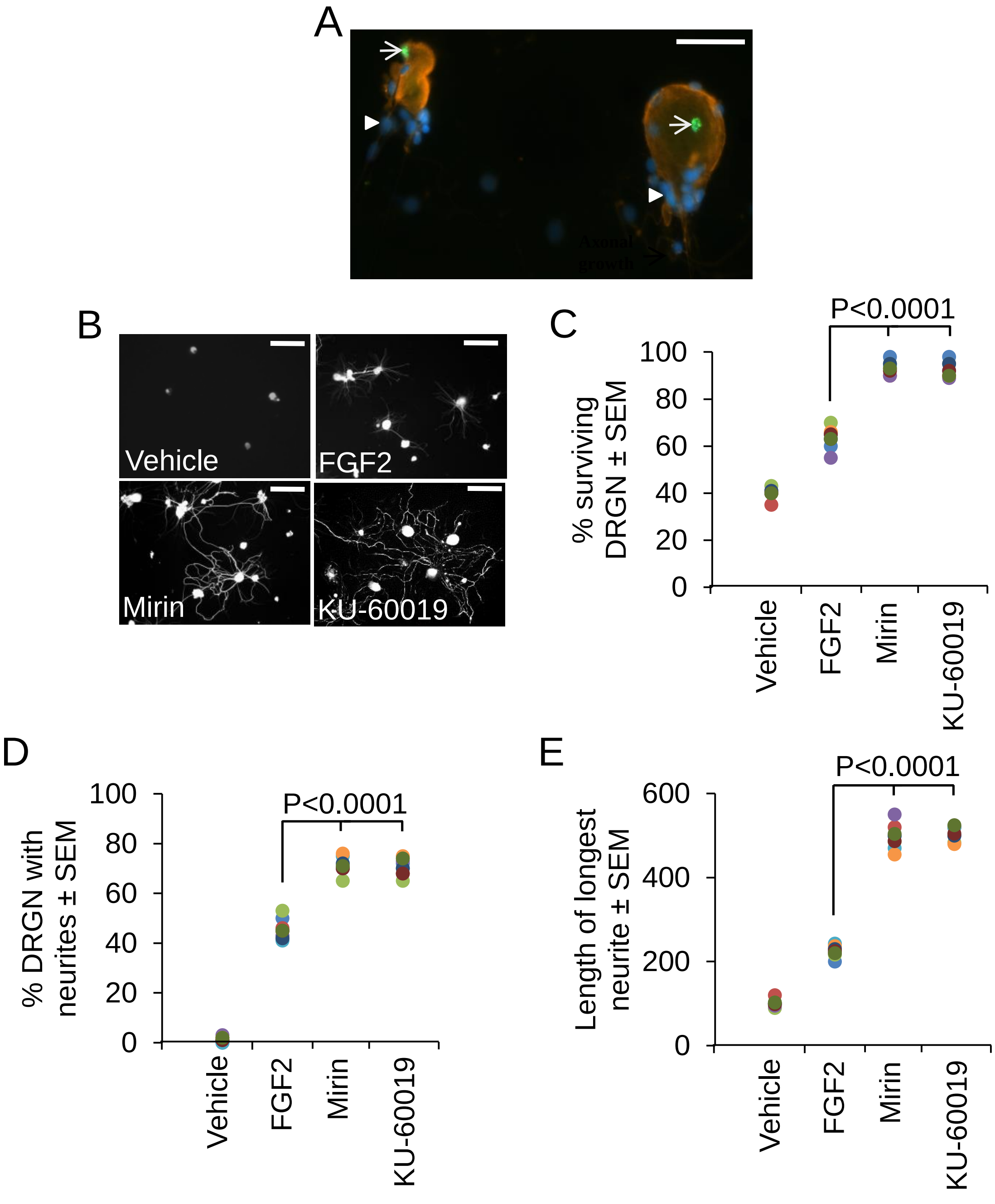


**Supplementary Fig. 3**

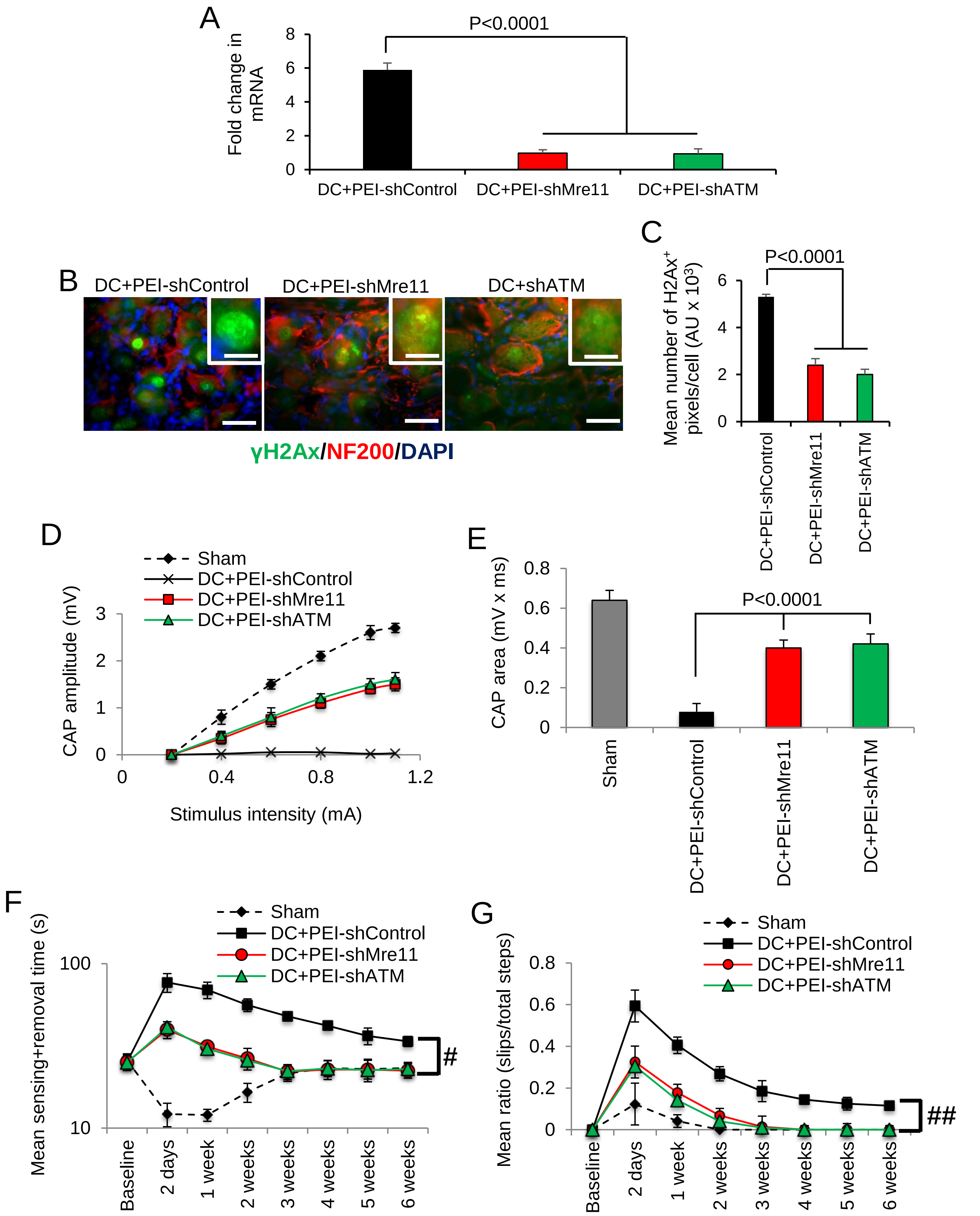


**Supplementary Fig. 4**

**Supplementary Table 1.** Extended Statistics to show degrees of freedom (df), F and actual P values.

| **Figure** | **Exp. unit** | **Degrees of freedom** | **F** | **Actual P value** |
|---|---|---|---|---|
| Fig. 1B | Fly | 15 | 51.3 | 0.0001 |
| Fig. 1C | Fly | 3, 586 | 36.8 | 0.0001 |
| Fig. 1E | Fly | 4, 122 | 15.9 | 0.0001 |
| Fig. 2A | Fly | 3 | 29.1 | 0.0001 |
| Fig. 2B (PDF+) | Fly | 2 | 46.0 | 0.0001 |
| Fig. 2B (PDF-) | Fly | 2 | 42.1 | 0.0001 |
| Fig. 2D | Fly | 3 | 14.5 | 0.0023 |
| Fig. 3B | Blots | 3 | 2596.7 | 0.0001 |
| Fig. 3C | Wells | 3 | 467.3 | 0.0001 |
| Fig. 3E | Wells | 3 | 2984.3 | 0.0001 |
| Fig. 3F | Wells | 3 | 1029.5 | 0.0001 |
| Fig. 4B | Blots | 2 | 1790.8 | 0.0001 |
| Fig. 4D | Blots | 2 | 421.0 | 0.0001 |
| Fig. 4F | Retinae | 3 | 2118.9 | 0.0001 |
| Fig. 4H | Nerves | 2 | 5430.0 | 0.0001 |
| Fig. 5C | Rats | 2 | 702.3 | 0.0001 |
| Fig. 5E | Blots | 2 | 198.4 | 0.0001 |
| Fig. 6B | Rats | 2 | 780.0 | 0.0001 |
| Fig. 7C | Rats | 3 | 3023.1 | 0.0001 |
| Supplementary Fig. 2B | Fly | 3, 586 | 2.842 | 0.0372 |
| Supplementary Fig. 2E | Fly | 3 | 28.70 | 0.0001 |
| Supplementary Fig. 3C | Wells | 3 | 623.7 | 0.0001 |
| Supplementary Fig. 3D | Wells | 3 | 1063.8 | 0.0001 |
| Supplementary Fig. 3E | Wells | 3 | 1159.9 | 0.0001 |
| Supplementary Fig. 4A | Rats | 2 | 812.7 | 0.0001 |
| Supplementary Fig. 4C | Rats | 2 | 407.9 | 0.0001 |
| Supplementary Fig. 4E | Rats | 3 | 648.1 | 0.0001 |

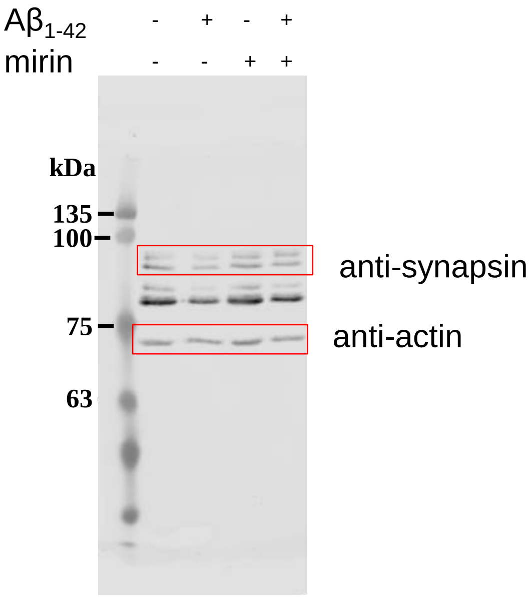


### Fig. 2D Original blot

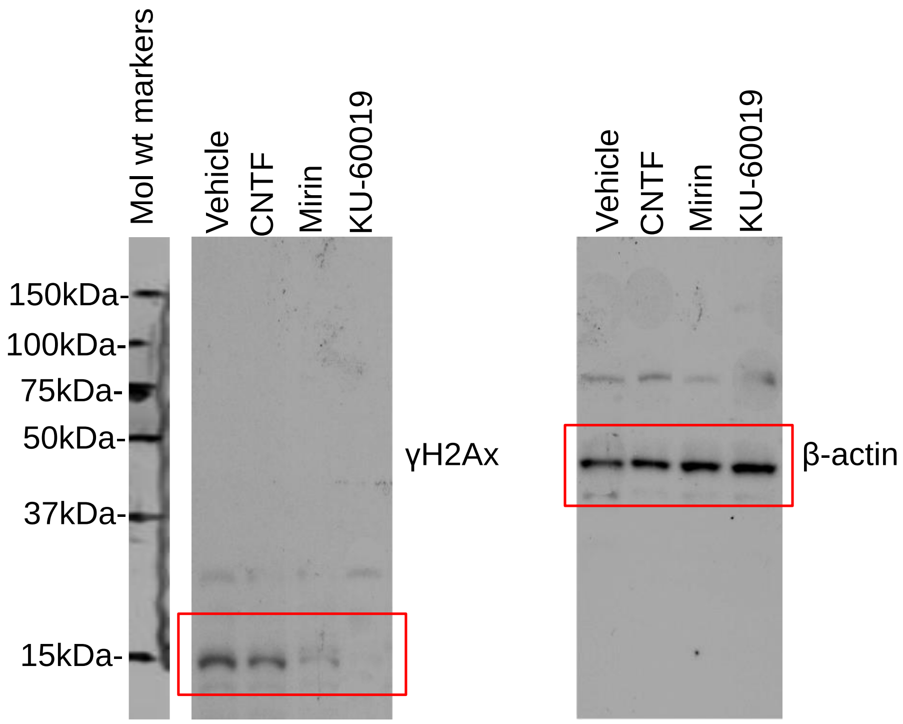


**Fig. 3A Original blot**

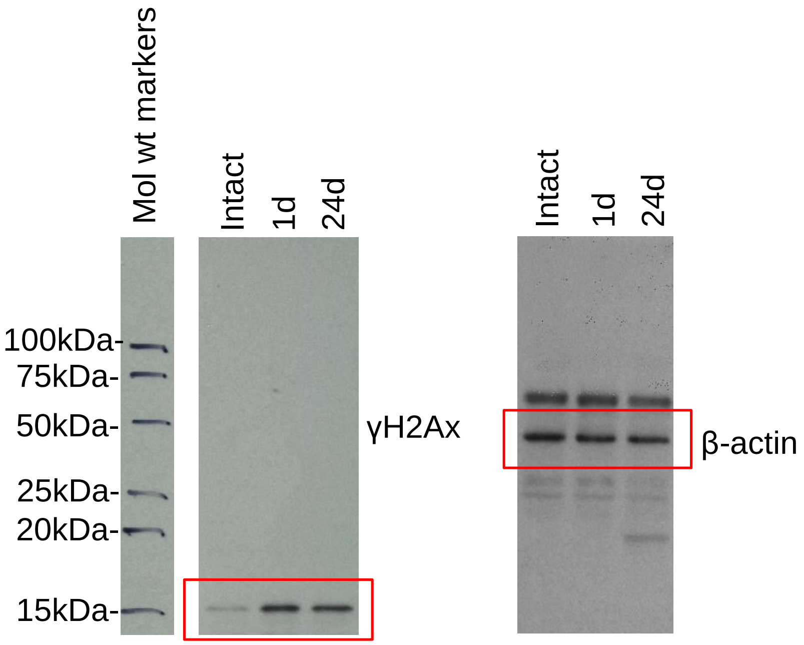


**Fig. 4A Original blot**

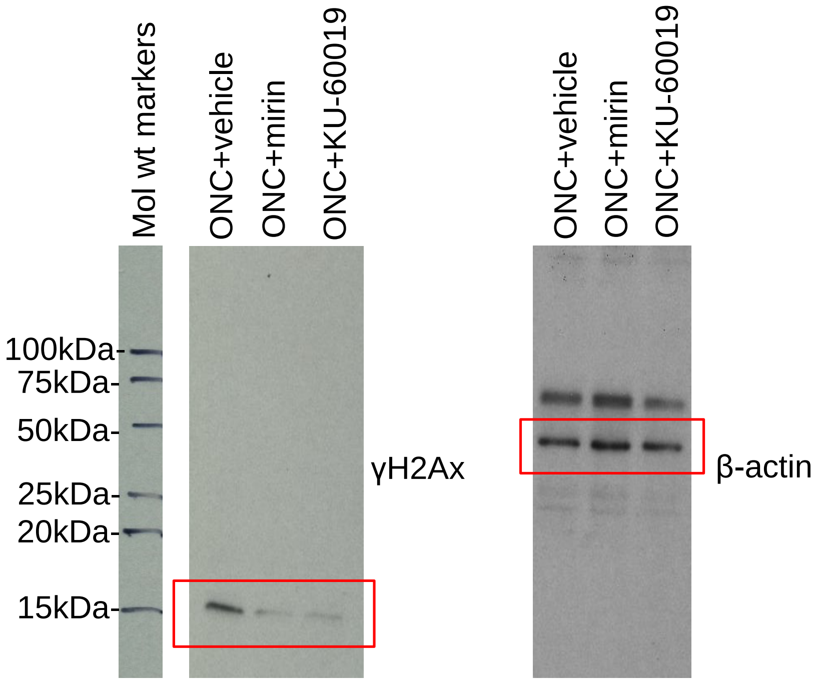


**Fig. 4C Original blot**

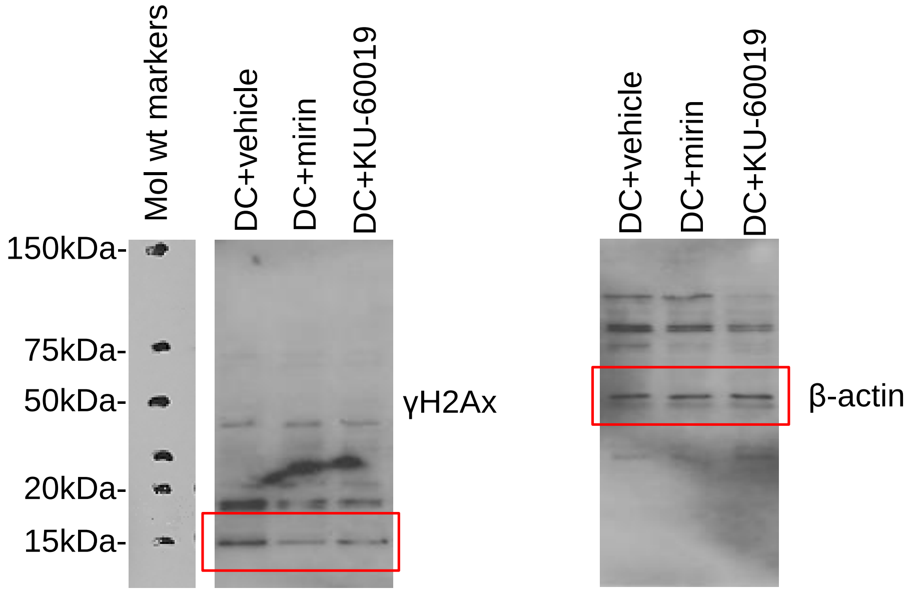


**Fig. 5D Original blot**

Originals of the western blots used for quantification of pTau from *Drosophila* heads shown in Supplementary figure S2.

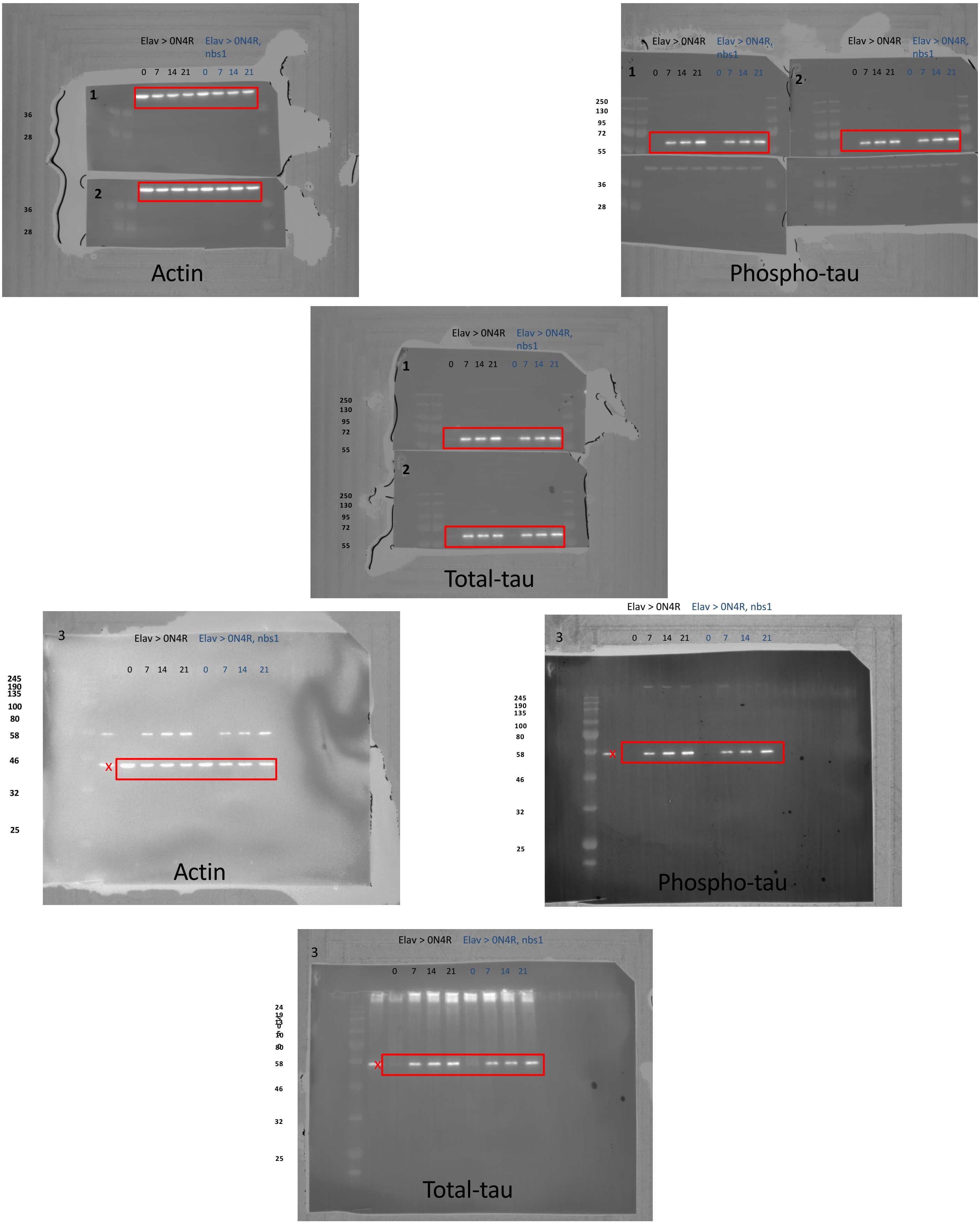
